# Modeling the effects of EMT-immune dynamics on epithelial cancer progression

**DOI:** 10.1101/615971

**Authors:** Daniel R. Bergman, Matthew K. Karikomi, Min Yu, Qing Nie, Adam L. MacLean

**Affiliations:** Department of Mathematics, University of California, Irvine, Irvine, CA 92697; Department of Stem Cell & Regenerative Medicine, Keck School of Medicine, University of Southern California, Los Angeles, CA 90033; Department of Cell and Developmental Biology, University of California, Irvine, Irvine, CA 92697; Department of Biological Sciences, University of Southern California, Los Angeles, CA 90089

## Abstract

**D**uring progression from carcinoma in situ to an invasive tumor, the immune system is engaged in complex sets of interactions with various tumor cells. Tumor cell plasticity also alters disease trajectories via epithelial-to-mesenchymal transition (EMT). Several of the same pathways that regulate EMT are involved in tumor-immune interactions, yet little is known about the mechanisms and consequences of crosstalk between these regulatory processes. Here we introduce a multiscale evolutionary model to describe tumor-immune-EMT interactions and their impact on epithelial cancer progression from in situ to invasive disease. Through in silico analyses of large patient cohorts, we find controllable regions that maximize invasion-free survival. We identify that delaying tumor progression depends crucially on properties of the mesenchymal tumor cell phenotype: its growth rate and its immune-evasiveness. Through analysis of EMT-inflammation-associated data from The Cancer Genome Atlas, we find that association with EMT significantly worsens invasion-free survival probabilities in support of our model, and we predict new genes influencing outcomes in bladder and uterine cancer, including FGF pathway members. These results offer novel means to delay disease progression by regulating properties of EMT through specific gene interactions, and demonstrate the importance of studying cancer-immune interactions in light of EMT.

## 1 Introduction

The majority of deaths from cancer are due to metastasis of the disease [1]. It is thus of critical importance to understand better the progression from in situ to invasive disease. Underlying this progression are genetic and epigenetic events, including mutations in pathways critical to the success of the cancer cell (driver mutations) [2]. These pathways include cell proliferation, apoptosis, and immunogenicity.

Cancer and the immune system interact in myriad ways. The immune system modulates the tumor microenvironment (TME), since immune signals that affect the tumor can be amplified or repressed through feedback in response to local inflammatory signals. This complex cell signaling occurs alongside the targeting (and potential eradication) of the tumor by immune cells [3].

The effects of the immune system on a tumor can be broadly summarized into two branches. The cytotoxic branch of the immune system, such as natural killer cells (NKs) and cytotoxic T cells (CTLs), seek out and lyse tumor cells. Upon carrying out their effector functions, these cytotoxic cells lose efficacy or deactivate [4]. The regulatory branch of the immune system (Tregs, and other factors), inhibits the effective functioning of the cytotoxic branch [5]. Inflammation can increase the probability of cancer incidence and progression, with some of the most pronounced effects seen for tumors originating in gastrointestinal and pancreatic tissues [6, 7]. Recent work has shown, contrary to the typical effects of inflammation on cancer, that under certain conditions inflammation may not be oncogenic but rather onco-protective [8].

Immunotherapies are beginning to realize their potential, with significant impacts on patient health and survival [9, 10], and may even provide a cure for certain hematopoietic cancers via anti-CD19 CAR-T cells [11]. The presentation of antigens on tumor cells is recognized by innate immune cells that are transported to lymph nodes where T cells (and other components) can be activated [12]. The tumor also engages in processes that can indirectly modify the TME, for example by releasing transforming growth factor beta (*TGF-β*), which can shift the TME towards a tumor-supportive environment by enhancing immunosuppression via activation of Tregs [12].

Epithelial-to-mesenchymal transition (EMT) describes a reversible process by which cells displaying an epithelial phenotype transition into cells with a mesenchymal phenotype. Epithelial cells are – in part – defined by tight cell-cell adhesion. Mesenchymal cells exhibit less adhesion, greater ranges of motility, and may possess stem-like properties [13], although controversy regarding ‘stemness’ and EMT remains [14, 15]. Recent work has shown that – rather than being a binary process – at least two stable intermediate EMT states exist [16, 17]. Ongoing investigations into the plasticity and stability of EMT overlap with discussions elsewhere, e.g. of discrete vs. continuous processes during cell differentiation [18]. Intermediate states have emerged as a central mechanism by which cell fates (and the noise inherent within them) can be controlled [19–21].

Two features of the mesenchymal phenotype are of particular relevance in the context of cancer-immune interactions. i) mesenchymal tumor cells proliferate less than epithelial cells, we refer to this as mesenchymal growth arrest (MGA), and can be considered related to (in the sense of quiescence) the “stemness” phenotype of the mesenchymal tumor cells [22]. ii) mesenchymal cells are less susceptible to immune clearance [23]. As a cell is targeted by cytotoxic immune cells for clearance, a physical connection between the two cells must be established. This immunological synapse – mediated in part by T-cell receptors bound to antigens and the major histocompatibility complex on the target cell – is down-regulated in mesenchymal cells, thus inhibiting formation of the synapse [23]. We refer to this phenotype as mesenchymal immune evasion (MIE).

In addition to the prominent role it plays in metastasis, EMT has more recently been shown to also regulate other aspects of tumor progression [13, 24] and tumor dormancy [25]. *TGF-β*, a master regulator of EMT [26], is at once implicated heavily in tumor-mediated immune responses, since Tregs release *TGF-β* upon arriving at the tumor site [23]. In hepatocellular carcinoma, for example, there is direct evidence linking Treg-secreted *TGF-β* with EMT [27]. Thus, even by considering only the *TGF-β* pathway, we find compelling evidence that these three core components (the tumor, the immune system, and EMT) all interact. It therefore strikes us as a priority to develop models to understand how the interactions between each of these three components affect cancer incidence and progression.

Mathematical oncology, that is, mathematical models of cancer incidence, progression, and treatment, has become a well-developed field; many models have offered insight into the cellular interactions underlying cancer and its interplay with the immune system, including older [28–30] and more recent works [31–44]. These studies have increased our understanding of how tumors grow in the presence of various immune components, and how treatment regimes can be designed to maximize the efficacy of cytotoxicity while minimizing risks to the patient. However, to our knowledge no models have addressed how the effects of EMT alter interactions between the immune system and cancer, and the subsequent implications for treatment.

Here we develop a model with the goal of studying interactions between the tumor, the immune system and EMT. We seek to describe a set of crucial molecular and cellular interactions in epithelial tumor cells, including effects due to DNA damage and mutation, to investigate the probability that in situ tumors will progress and, if so, when. A recent model of cancer-immune interactions [8] described the effects of the TME on the risk of cancer, and we build on the core cell cycle component of this model, adding significant new interactions to the immune component of the model (which was previously modeled by a single interaction), as well as adding the effects of EMT. In doing so, we shift the focus of the previous model from cancer initiation to cancer progression. We do this to reflect the fact that cancer progression hinges on escape from the immune system and the fact that EMT has a more well-defined role during progression and metastasis. We seek to understand whether this more complex immune module will change our understanding of inflammatory effects on the tumor, and how the epithelial-mesenchymal axis influences these.

In the next section we develop the model, explaining the intuition behind each of its components. We go on to analyze its behavior: global “*one-at-a-time*” sensitivity analysis identifies parameters that are crucial for progression. We study these in more depth, focusing on the competing effects of EMT and of the immune system on progression, and discover that EMT intricately regulates progression: under certain regimes a careful balance of EMT- and immune-driven processes can significantly prolong invasion-free survival. To test these predictions, we analyze data from the Cancer Genome Atlas (TCGA) using a new pipeline, and find strong evidence for the synergistic effects of inflammation and EMT, predicted through co-expression effects, for patients with bladder and uterine cancers.

#### Quick Guide to Equations and Assumptions

To become invasive, An in situ tumor relies on mutations to alter cellular signaling pathways that enable cancer progression. The immune system simultaneously responds to the tumor upon recognition of neoantigens, and shapes the TME through dynamic inflammatory and regulatory signals.

To capture these dynamics, we developed a non-spatial agent-based model. Tumor cells are modeled individually as agents and immune cell populations are described homogeneously by differential equations. We consider two tumor cell types: epithelial tumor cells (ETCs) and mesenchymal tumor cells (MTCs). Time is treated discretely in 18 hour steps; approximating the time of one cell cycle. During each cell cycle, tumor cells can either undergo division, apoptosis, immune clearance, or arrest in G_0_. The likelihood that a cell will proliferate depends on the tumor size (competition for resources) and on cell-intrinsic factors. The likelihood that a cell will undergo apoptosis is constant but varies between cell types (ETC and MTC). The likelihood of immune clearance depends on the number and type of mutated cells in the tumor, cytotoxicity, regulatory cells, and cell-intrinsic factors. As tumor cells proliferate, DNA damage can occur, and over time they become increasingly likely to acquire pathway mutations that change their propensities for proliferation or cell death. Natural killer (NK) cells identify and clear tumor cells, a process which results in neoantigens priming and activating T cells in local lymph nodes. T cells can subsequently infiltrate the TME. At the tumor site, cytotoxic T cells (CTLs) lyse tumor cells, and T regulatory cells (Tregs) suppress cytotoxic activity. The following equation determines probability (per cell cycle) that a tumor cell will be lysed by a CTL:

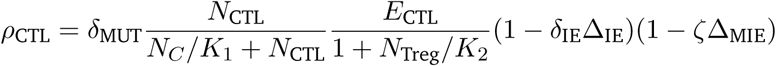

Here, *δ*_MUT_ is 1 or 0 depending on whether or not the cell has a mutation. The second term is a hill function modeled after [45] and the third term describes onco-protective effects of Tregs. The second-to-last term describes the increased immune-evasiveness that can occur following mutation in the immune evasion pathway, and the last term quantifies the additional immune evasiveness of associated with MTCs. The inflammatory state of the TME thus impacts (through multiple factors) immune recruitment and cytotoxicity at the tumor site.

EMT impacts tumor cells through their proliferation and potential to evade the immune system. MTCs have reduced proliferation and increased immune evasiveness. *TGF-β*, an activator of EMT, is produced both by Tregs and tumor cells, thus connecting tumor-immune interactions with EMT, and as a result plays an important role in shaping tumor outcomes. To determine whether a cell undergoes EMT or MET, the *TGF-β* is randomly divided among the cells so that associated with each cell *i* is a value *τ*_*i*_. This value is given by the following equation:

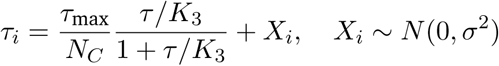

Here, *τ* is the concentration of *TGF-β* in the TME and *τ*_max_ represents a limit on the amount of *TGF-β* that can be absorbed by all cells; a Gaussian noise term is added. For each cell, *τ*_*i*_ is summed with the current EMT score for that cell and if the result is above a threshold value, the cell undergoes EMT, otherwise MET. This summation expresses the assumption that the epithelial and mesenchymal phenotypes are stable and cells will move along the EMT spectrum towards the equilibrium they identify as under typical conditions. Our measured outcome is the time at which the cancer becomes invasive, determined through the proportion of tumor cells harboring mutations in pathways that permit escape, relative to the total tumor cell population.

## 2 Methods

Here we briefly describe to core components of the model. Full details and equations are provided in the Supplementary Data. We develop an agent-based model to describe the relationships between cancer, the immune system, and EMT, building on the cell-cycle and tissue-cell components described in [8]. The agents in the model are the cells that have already formed an in situ tumor yet lack key pathway mutations to become invasive. In the process of the simulation, these cells can acquire mutations altering any of three key pathways (Fig. 1A).

**Figure 1:**
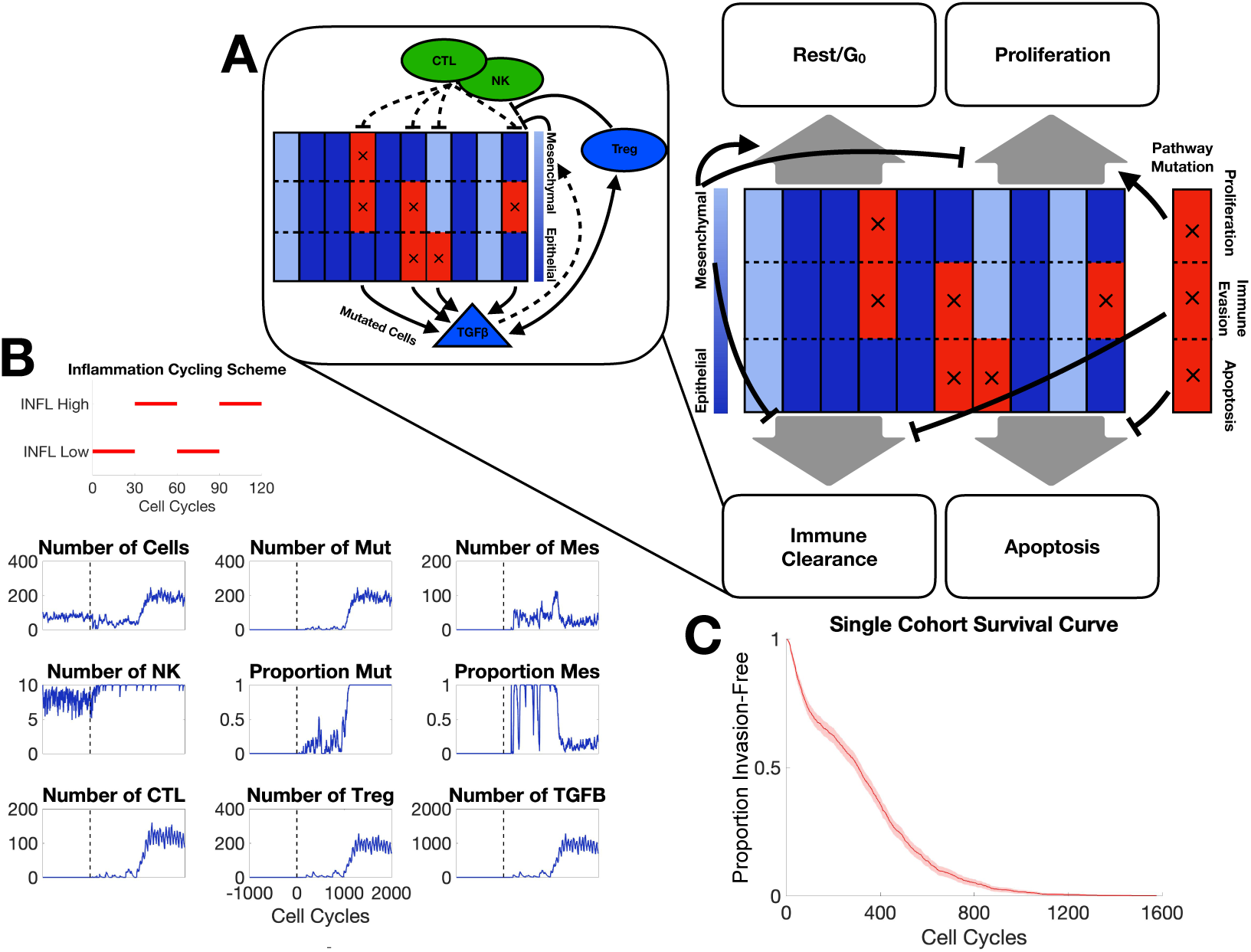
**A.** Schematic depiction of agent-based model components; each of the 10 columns represents one tumor cell divided into three compartments representing the state of three pathways with tumorigenic potential; blue/red denote baseline/altered pathway activity. Black arrows depict cell fate regulation in each cell cycle. Inset depicts major interactions between the immune system and tumor cells. **B.** A representative simulation of one patient. The parameter values used can be found in Table S2. The inflammation cycling scheme (red) is shown above the patient dynamics. The vertical dashed line denotes the end of the warmup period. Mut: malignant cells; Mes: mesenchymal cells. **C.** Survival curve for one cohort of patients for parameter values given in Table S2.

We model immune cells as continuous variables, i.e. we assume that the tumor microenvironment is well-mixed with regards to the infiltrating immune cells. The cytokine *TGF-β* is also assumed to be well-mixed in the tumor microenvironment. Tumor cells can take on either epithelial or mesenchymal phenotypes in a plastic manner: these phenotypes depend on both the TME and cell-intrinsic factors. While the EMT score is continuous, a threshold determines if a given cell is labeled as epithelial or mesenchymal (Fig. 1).

### 2.1 Tumor evolution

Associated with each tumor cells are two essential features: their mutational signature and their EMT score. We consider three idealized pathways that can be mutated: proliferation, when altered this increases the probability of the cell proliferating within each cell cycle; apoptosis, when altered this decreases the probability of a cell undergoing apoptosis; and immune evasion, when altered this decreases the probability that a mutated cell will be cleared by immune components.

### 2.2 Immune population dynamics

The immune system is modeled by three immune cell types: NKs, CTLs, and Tregs. The NKs and CTLs act on the system by recognizing invasive cells and clearing them. Upon clearance, they are deactivated and removed from the immune population. Tregs suppress the function of NKs and CTLs (reduce tumor cell clearance), and in addition, release *TGF-β* which further shapes the TME by pushing tissues cells more towards a mesenchymal phenotype.

### 2.3 Periodic cycling inflammation states

Inflammation is modeled as a cycling scheme between low and high inflammatory states, with varying on/off durations and intensities. For the purpose of simulation we consider the default state to be low inflammation, and update the immune activity parameters whenever a switch to the high state occurs. In Supplemental Table S2 we give full details of parameter settings during low and high inflammation.

### 2.4 Epithelial-to-mesenchymal transition

Each tissue cell has an EMT score between 0 and 1 with is set according to the concentration of *TGF-β* in the TME. Above a threshold, the cell acquires the phenotype of a mesenchymal tumor cell (MTC); otherwise, it is an epithelial tumor cell (ETC). For the purpose of simulation, ETCs are considered to be in the base state, and MTCs will have a subset of their parameters updated. In modeling EMT this way, we are assuming that the same factor, *TGF-β*, drives EMT both at initiation and through progression of cancer.

Cells that have undergone EMT (i.e. MTCs) experience a reduction in proliferation, referred to as mesenchymal growth arrest (MGA), and a decrease in the likelihood that they will be cleared by immune cells (NKs or CTLs), referred to as mesenchymal immune evasion (MIE). Both these parameters lie within the range [0, 1], thus we can sufficiently sample from their joint parameter space to explore it in depth with the need for informative priors to constrain their values.

### 2.5 Model simulation

#### Initial conditions

Simulations are initialized with *N*_0_ in situ tumor cells, determined by the choice of parameter values. A number of warmup cycles are run so that the model reaches steady state. During warmup, no mutations occur, and the only immune cells present are NKs. After warmup, mutations are permitted. Cells that do not mutate undergo an increase in their probability of mutation in a later cell cycle.

#### Tumor cell fate

During each cell cycle, the fate of each cell is assigned: proliferation, apoptosis, immune clearance, and rest in *G*_0_, according the model rules. The probability of proliferation is affected by mutations to the proliferation pathway (increased) and my cells in a mesenchymal state (decreased). Probabilities of immune clearance are affected by the number of mutations harbored: cells with more mutations are assumed to be more immunogenic and have a higher probability of being cleared by the immune system, unless the cell has a mutation in the immune evasion pathway. Cells in a mesenchymal state can exhibit greater capacity to evade immune clearance.

#### Completing the cell cycle

Once all tumor cells have been updated and fates chosen accordingly, non-tumor model components are updated. Immune cell populations are updated in two steps. First, immune cell exhaustion is calculated based on the number of tumor cells cleared, e.g. clearance of one tumor cell by an NK cells results in the NK cell population decreasing by one. Second, all immune cells (NKs, CTLs, Tregs) are updated according to a system of coupled ordinary differential equations that govern their population dynamics. CTL and Treg recruitment rates are dependent on the number of tumor cells; in addition *TGF-β* enhances the recruitment rate of Tregs.

At the end of each cell cycle, new mutations can occur in cells that have undergone division, according to cell-specific probabilities that increase if no mutation occurs are reset to 0 in the event of a mutation. Finally, the concentration of *TGF-β* and the EMT score for each cell are updated. Tregs and (to a lesser extent) invasive tumor cells are the sources of *TGF-β*; the total concentration per cell cycle is divided randomly among tumor cells. EMT is then assessed, depending on the EMT score of the cell and the local concentration of *TGF-β*.

#### Mutational burden and progression to invasive disease

At the end of each cell cycle, the proportion of tumor cells that are invasive is calculated based on their mutational burden, and if it is above a certain threshold, the tumor is declared to have progressed to an invasive state and the simulation ends. The time to invasion is calculated as the time from the start of the simulation, minus the warmup period, until the invasive state is reached. Simulations run until either this occurs or until the maximum number of cell cycles has been reached.

### 2.6 Parameter estimation and sensitivity analysis

To study parameter sensitivity, we implemented Morris a one-step-at-a-time global sensitivity analysis. Parameters are varied one at a time from a set of sampled “base” points and the resulting simulations recorded [46,47]. For each run we simulated 1000 patients, and initialized the Morris sampling with 30 points in parameter space (at least 10 are recommended in [47]. Parameter sampling a choice of prior parameter distributions. For many parameters, such as for the immune population dynamics, measurements or estimates were available from literature [45]. For parameters such as MIE and MGA related to the mesenchymal phenotype, little prior information was available, thus these was sampled across all possible values in [0, 1]. Tumor size in the model was scaled from cell numbers on the order of 109 cells [45] to the order of 102, and parameter values were scaled accordingly. Where parameter estimates existed, the prior for parameter *θ*_*i*_ is given as *θ*_*i*_ ∼ N(*m*_*e*_, 2*m*_*e*_), where *m*_*e*_ is the previous estimate and we take twice this value as the variance to obtain a range of samples that does not rely too heavily on previous work. The Morris algorithm computes the sensitivity, *µ*^*^, as the average of the absolute change of the output, which in our model is the area under the survival curve (Fig. 2).

**Figure 2:**
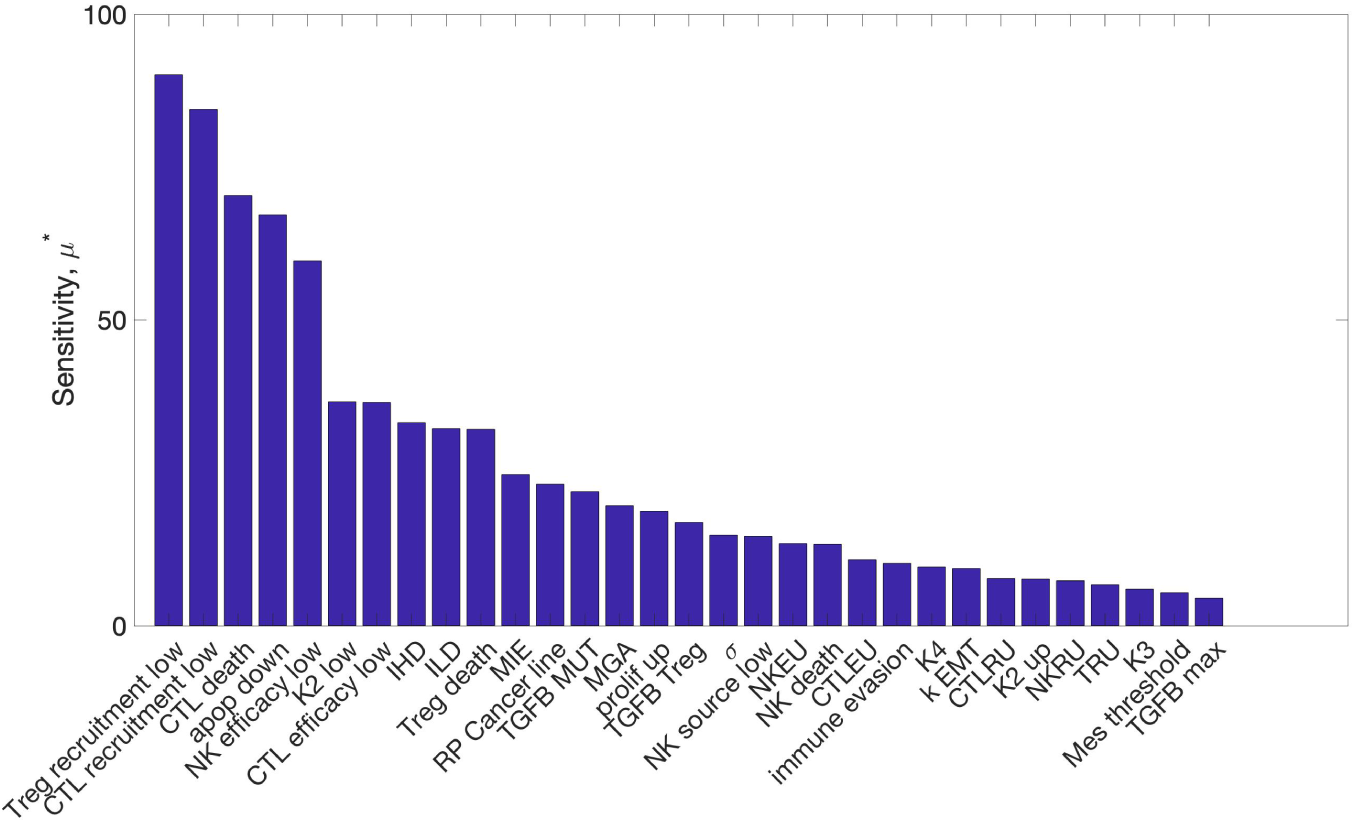
Global sensitivity analysis of model parameters. The sensitivity (*µ*^*^) denotes the average absolute change in the time to invasion over the range of variation of the parameter.

### 2.7 Analysis of patient survival data from TCGA

We obtained primary tumor bulk mRNA sequencing and censored survival data for individuals monitored by cancer type from the Cancer Genome Atlas (TCGA) [48,49], accessed through the Genomic Data Commons portal [50]. We developed methods to study: i) how the synergistic effects of EMT + inflammation compare to the effects of each of these individually; and ii) the importance of mesenchymal proliferation rates in determining cancer prognosis (Supplemental Figure S5), which allow us to test predictions from the agent-based model.

For (i), we identified cases where synergistic effects due to the combination of EMT and inflammation pathways have greater influence on overall survival than the effects of either of these factors alone. For each cancer type, we obtained from MSigDB [51] gene sets that contain EMT or Inflammation-related genes and, for each gene set, we tested whether EMT, inflammation, or the combination of these two effects best predicts overall survival. We selected for those gene sets that exhibit strong synergistic effects as identified by a Cox proportional-hazard (CPH) model. For tumor types where synergistic effects were most evident, we asked whether unsupervised clustering of patients, based on a low-dimensional representation of the combination gene set, could predict statistically significant differences in overall survival via the Kaplan-Meier (KM) model. For tumor types where the both the CPH and KM analysis were consistent, we conducted further analysis (ii) of the role played by proliferation on tumor invasiveness.

**Table 1:**
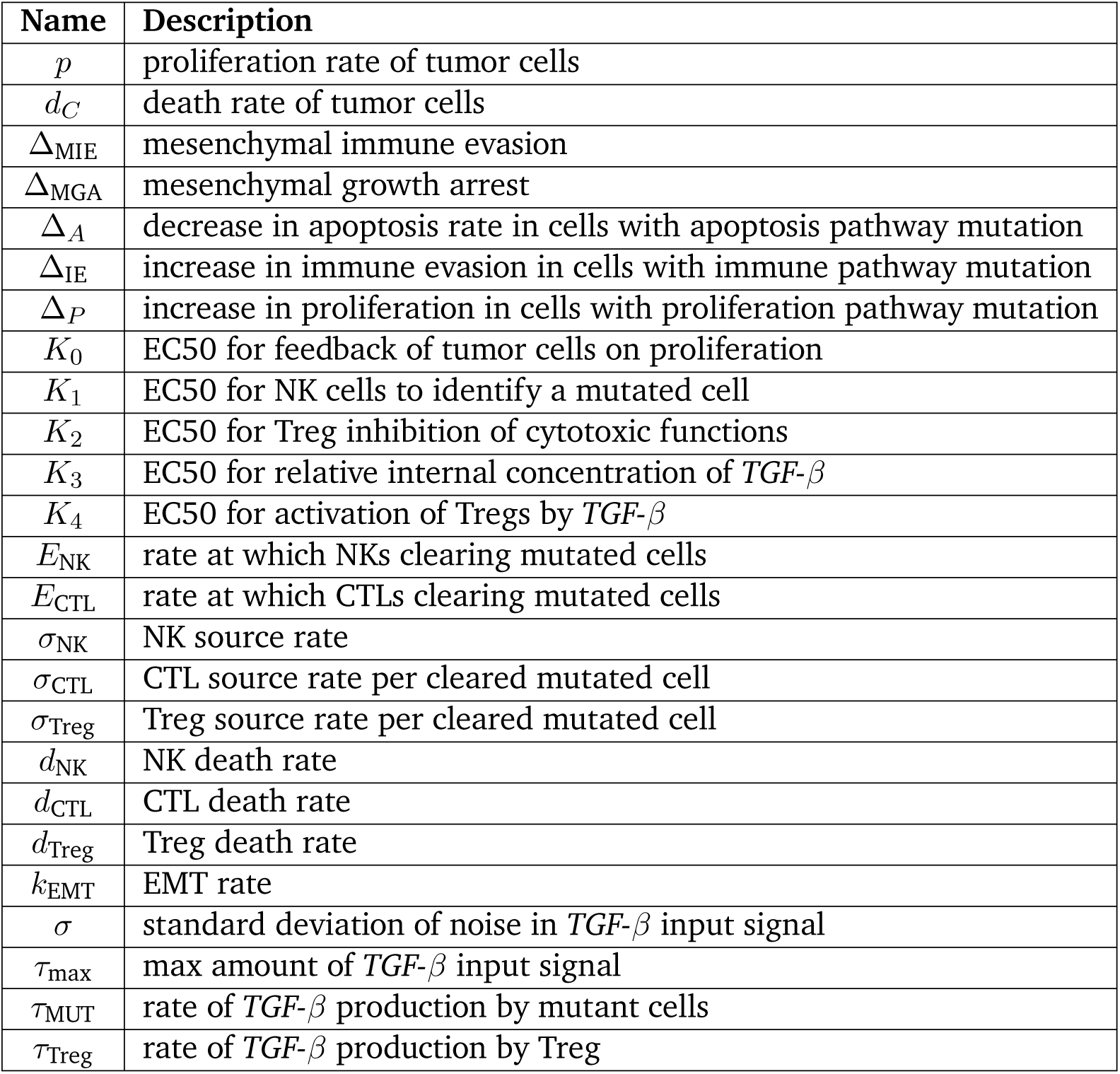
Description of key model parameters. Note that some are not constant as they can be affected by the inflammation state of the system.

| Name | Description |
| --- | --- |
| $p$ | proliferation rate of tumor cells |
| $d_C$ | death rate of tumor cells |
| $\Delta_{\text{MIE}}$ | mesenchymal immune evasion |
| $\Delta_{\text{MGA}}$ | mesenchymal growth arrest |
| $\Delta_A$ | decrease in apoptosis rate in cells with apoptosis pathway mutation |
| $\Delta_{\text{IE}}$ | increase in immune evasion in cells with immune pathway mutation |
| $\Delta_P$ | increase in proliferation in cells with proliferation pathway mutation |
| $K_0$ | EC50 for feedback of tumor cells on proliferation |
| $K_1$ | EC50 for NK cells to identify a mutated cell |
| $K_2$ | EC50 for Treg inhibition of cytotoxic functions |
| $K_3$ | EC50 for relative internal concentration of $TGF-\beta$ |
| $K_4$ | EC50 for activation of Tregs by $TGF-\beta$ |
| $E_{\text{NK}}$ | rate at which NKs clearing mutated cells |
| $E_{\text{CTL}}$ | rate at which CTLs clearing mutated cells |
| $\sigma_{\text{NK}}$ | NK source rate |
| $\sigma_{\text{CTL}}$ | CTL source rate per cleared mutated cell |
| $\sigma_{\text{Treg}}$ | Treg source rate per cleared mutated cell |
| $d_{\text{NK}}$ | NK death rate |
| $d_{\text{CTL}}$ | CTL death rate |
| $d_{\text{Treg}}$ | Treg death rate |
| $k_{\text{EMT}}$ | EMT rate |
| $\sigma$ | standard deviation of noise in $TGF-\beta$ input signal |
| $\tau_{\text{max}}$ | max amount of $TGF-\beta$ input signal |
| $\tau_{\text{MUT}}$ | rate of $TGF-\beta$ production by mutant cells |
| $\tau_{\text{Treg}}$ | rate of $TGF-\beta$ production by Treg |

For (ii), we utilized the TCGA outcome “disease-free interval” (DFI) as it best resembles the invasion-free survival metric used in modeling (in many cases, disease may be undetected until it becomes invasive). We noticed that these times were highly bi-modal, suggesting that the GRN-based regulation of invasion could be learned via binary classification. We determined the patient DFI class by fitting these invasion times to a two-component Gaussian mixture model, which assigns each patient to either high-DFI or low-DFI. We then used Gaussian process classification to learn the regulatory structure of a group of mesenchymal proliferation genes based on their ability to predict DFI class. Specifically, we clustered the genes based on the rank order statistics of their respective maximum-a-posteriori (MAP) factor-analysis distances. Finally, we used simulations of the learned model to further examine the co-regulation of these genes, highlighting the interaction between immunity, tumor progression and invasiveness in the context of treatment-response. Full details of the methods used for this analysis can be found in the Supplementary Text.

## 3 Results

### 3.1 A multiscale agent-based model of EMT-immune-tumor cell interactions to study tumor progression

We begin by investigating general features of the model to establish baseline conditions and assess the impact of different model components on the key measured outcomes: the probability of progression, and the time to invasion. During the cell cycle, cell fate is determined by rules that are influenced by EMT and immune interactions (Fig. 1A), e.g. if a cell undergoes EMT, its probability of proliferation is reduced; if it gains a mutation in the apoptosis pathway, its probability of apoptosis is reduced. Meanwhile, NK cells and CTLs attempt to clear malignant tumor cells, and deactivate upon successful tumor cell clearance; Tregs inhibit this cytotoxic activity (1A Inset).

The inflammation cycling scheme for a typical in silico patient consists of alternating high and low regimes with corresponding effects on the cell populations (Fig. 1B). For this patient, after warmup, mutations are observed at a rate low enough that they are cleared by cytotoxic cells for about 700 cell cycles, after which the mutated and thus invasive cell population begins to grow, leading to large recruitment of CTLs and Tregs and a peak in the concentration of *TGF-β*. After 841 cell cycles, the proportion of invasive cells reaches 50%: the threshold defining progression, thus this patient has a time to invasion of 841 cell cycles, or 631 days. Beyond this timepoint, we see a rapid increase in the number of invasive cells until it comprises 100% of the tumor population. Interesting EMT dynamics are also observed, the proportion of MTCs peaks shortly after the tumor becomes invasive, subsequently the majority of cells transition back to an epithelial state. We observe that while the NK population varies little over the simulation, CTLs and Tregs both undergo large expansions. CTLs and Tregs also appear to oscillate, however note that this is a direct result of the inflammation state, and is not immune cell-intrinsic.

In order to quantify patient dynamics and invasion-free survival as a population level, we simulate large cohorts of patients similar to the single patient shown in Fig. 1B. For a cohort of 500 patients, we simulate survival curves and see that a large number progress quickly to form invasive tumors, whereas a few lie in the tail of the distribution after the mutagenic event that a large number of tumors quickly progress while others takes some time before progressing Fig. 1C. By approximately 1200 cell cycles (2.5 years), all tumors have become invasive..

### 3.2 Identification of key model parameters via global sensitivity analysis

Exploring the parameter spaces of systems biology models *adequately* is – in general – a hard problem. Fitting parameters via (Bayesian) parameter inference is advisable wherever possible [52]. Here, despite a wealth of data on tumor growth dynamics, a lack of sufficient molecular measurements (i.e. immune cell dynamics) precludes inference of the full model. In addition, while inference schemes for agent-based models are developing [53, 54], simulation times remain a hurdle [55]. Parameters for some components of the model studied previously can be constrained [8]. However, even here, new biological processes in the current system could push the model into new behavioral regimes. Thus to sample and characterize the parameter space of the model we use sensitivity analysis.

The results of Morris one-step-at-a-time sensitivity analysis on the 31 model parameters (Fig. 2) find a subset of parameters with much higher levels of sensitivity than others. The two most influential by this analysis are the recruitment rates of Tregs and CTLs in the low inflammation state. The parameters influencing EMT are also identified as influencing model outcomes. Since one goal of our analysis is to assess the specific effects of EMT on immune-cancer dynamics, parameters MIE and MGA are of particular interest. In addition, inflammation parameters controlling the periodic high/low inflammation states are of interest because they strongly influence model outcomes and are capable of being targeted by therapeutic treatments. For immune cell dynamics, the secretion of *TGF-β* by Tregs is found to be sensitive and thus will also be studied further below.

### 3.3 Mesenchymal properties dramatically alter invasion-free survival times

Mesenchymal tumor cells (MTCs) are characterized by changes in two parameters: mesenchymal immune evasion (MIE) and mesenchymal growth arrest (MGA). Here we assess the effects of each, alongside the effects of *TGF-β* through its production by Tregs. As MIE increases, the invasion-free survival decreases (Fig. 3A) for all sets of parameters studied: as the subpopulation of invasive cells becomes more resistant to immune clearance, the tumor as a whole grows more resilient and thus can grow faster (Fig. 3D).

**Figure 3:**
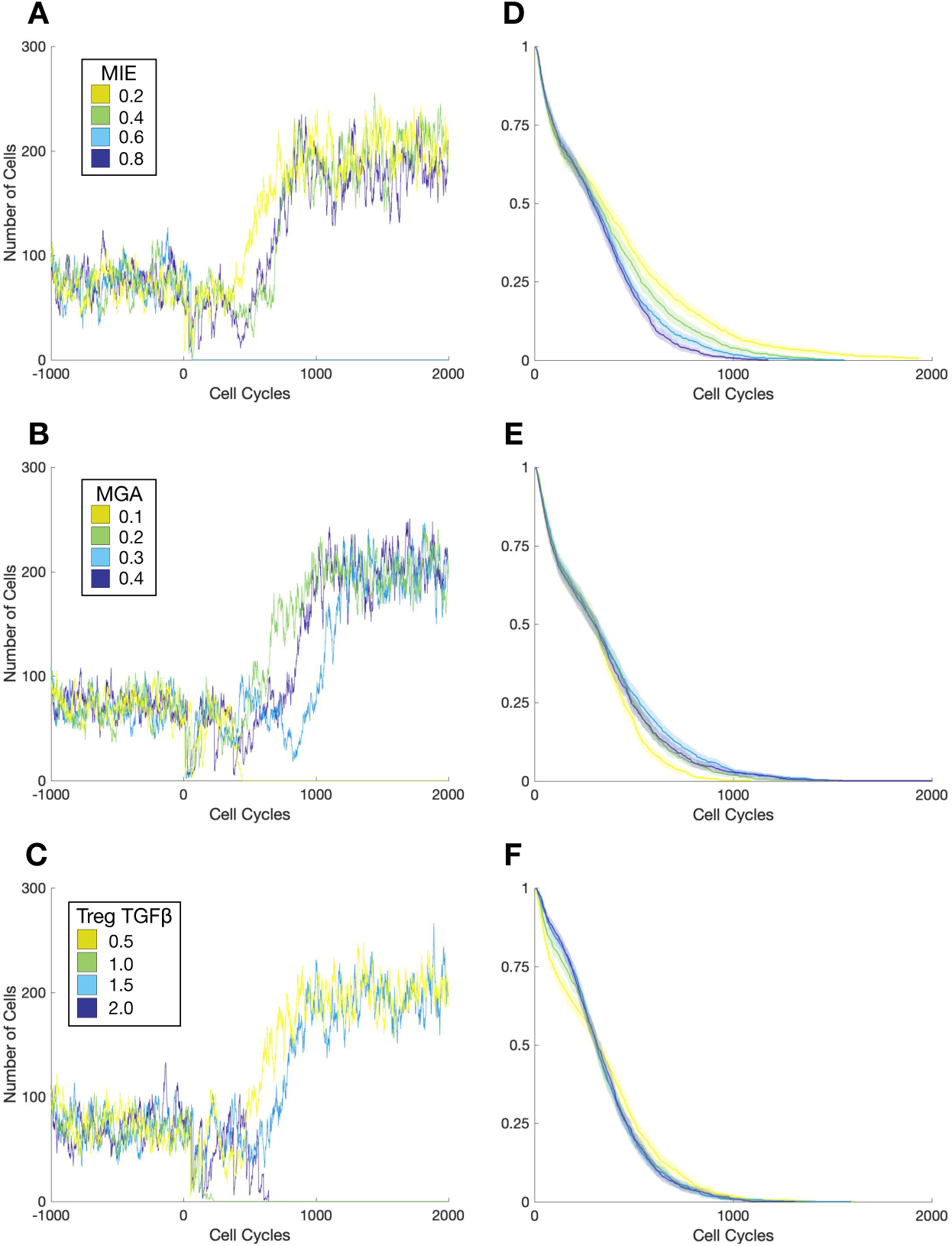
Effects of mesenchymal tumor cell properties on the time to invasion. Trajectories of one patient per cohort including warmup and 2000 cell cycles for **A.** mesenchymal immune evasion (MIE); **B.** mesenchymal growth arrest (MGA); **C.** Production of *TGF-β* by Tregs. **D.** Survival curve corresponding to changes in the parameter MIE (A) for a patient cohort of 1000. Shaded region represents the 95% confidence interval over the cohort. **E.** Survival curve corresponding to changes in MGA. **F.** Survival curve corresponding to changes in Treg production of *TGF-β*.

The relationship between MGA and invasion-free survival times displays a very different trend, and is non-monotonic with a local maximal value. For small values of MGA, increasing MGA results in increasing the invasion-free survival (Fig 3B, E). However for large values of MGA, invasion-free survival times decrease. This is explored further below.

*TGF-β* varies according to its production by tumor cells and its production by Tregs. We assess the effects of varying the production of *TGF-β* by Tregs on invasion-free survival (Fig 3C, F), and find that at lower production rates of *TGF-β*, the survival curve initially declines faster whereas higher production rates result in a steeper drop off in survival later. While lower values of *TGF-β* production lead to a steeper initial decline, these differences vanish for higher values of *TGF-β*. The steeper initial decline may be due to the rapid clearance of tumor cells by adaptive immune cells before Tregs have had sufficient time to modulate the TME (which they do through secretion of *TGF-β*).

### 3.4 A key EMT regime maximizes cancer-free survival time under chronic inflammation

To investigate how competing interactions within the inflammatory tumor microenvironment affect EMT, we explored the effects of varying inflammation on invasion-free survival. Patient cohorts were simulated under different inflammation regimes: permanently low inflammation; permanently high inflammation; or variable (periodic high/low) inflammation. Compared to the other inflammation states, permanently high inflammation results in outcomes that vary more subtly with changes in the mesenchymal parameters (Fig. 4). When the inflammation state is either permanently or temporarily low, surprising trends emerge. In both these cases, invasion-free survival time is negatively correlated with MIE, and a local maximum for the invasion-free survival time is found with respect to MGA (close to Δ_MGA_ = 0.2).

**Figure 4:**
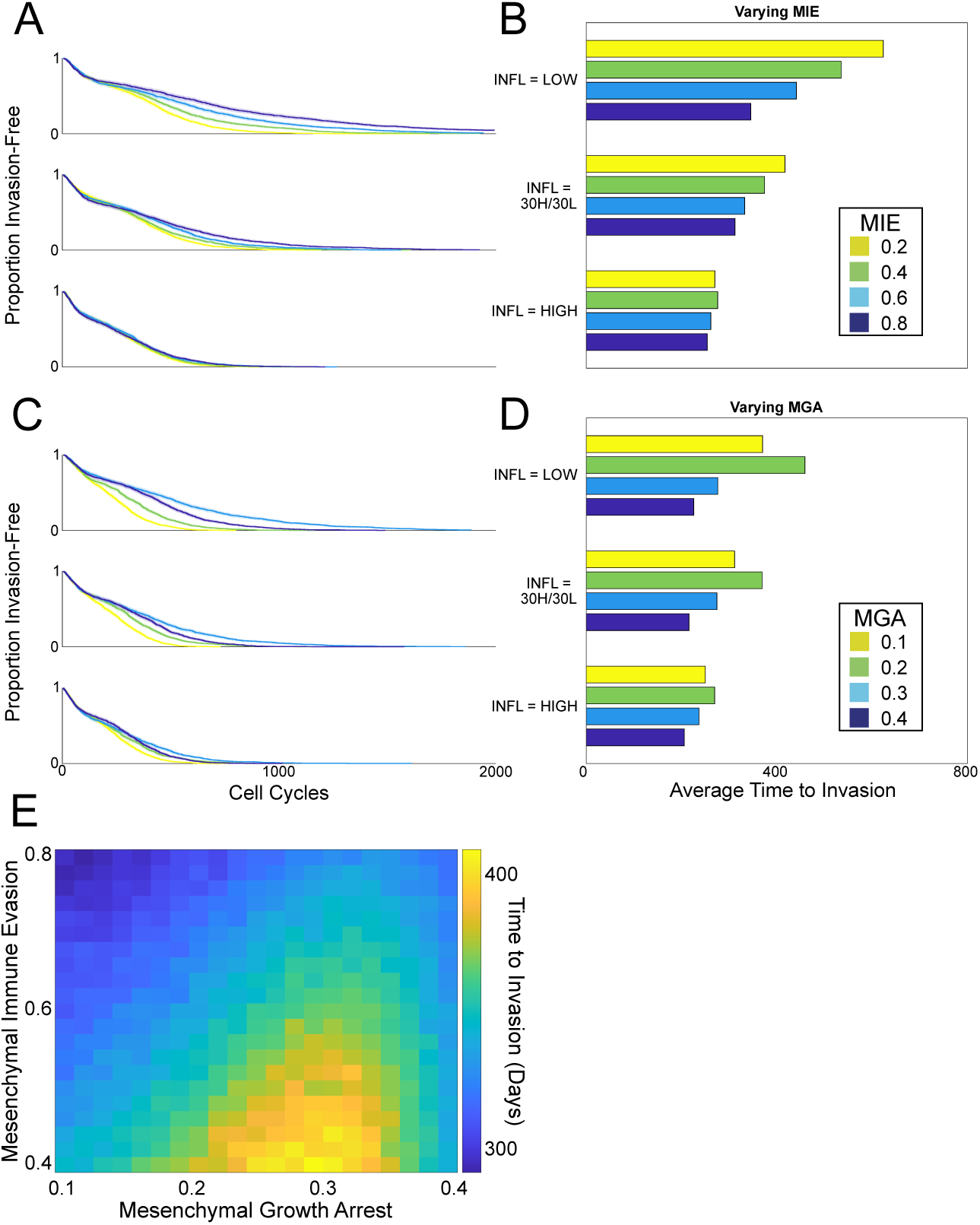
Effects of inflammation on the time to invasion under different cycling schemes. **A-B.** As MIE varies, survival curves (each of 200 patients) and corresponding bar plots to summarize the mean Time to Invasion for each cohort are shown. **C-D.** As MGA varies, survival curves and corresponding bar plots to summarize the mean Time to Invasion for each cohort are shown. **E.** Summary of the effects of MIE and MGA on invasion-free survival.

These differences in the mean invasion-free survival lead to striking variation in outcomes: tumors can be contained in situ for up to twice as long as they would otherwise be simply by varying the rates of mesenchymal growth arrest. These predictions point to intriguing therapeutic outcomes: a patient suffering intermittent high inflammatory attacks will benefit directly from EMT-directed therapies, however patients for whom a relatively high inflammation state is observed chronically will not obtain this benefit.

When MIE is varied under different inflammation cycling schemes, for all the periodic inflammation schemes studied, increasing MIE will decrease the invasion-free survival (i.e. worsen cancer progression and prognosis) (Fig. 4B). In the case of continuously high inflammation, the effects of MIE are minimal. Thus, under any inflammation regime with periods of low inflammation, as we might intuitively assume, any reduction in mesenchymal immune evasion will lead to improvements in patient outcomes. To summarize the mesenchymal properties of immune evasion and growth arrest, we plot the joint density of these parameters against the time to invasion (Fig. 4E): where we see that for a given value of mesenchymal immune evasion, there is a value of mesenchymal growth arrest that maximizes the time to invasion.

### 3.5 TCGA data analysis supports predictions highlighting importance of mesenchymal cell proliferation in determining outcome

Given the model prediction that mesenchymal proliferation rates exert essential control on invasion-free survival times (Fig. 4E), we performed analysis of 14 cancer types from TCGA to assess the importance of mesenchymal proliferation-associated genes using clinical outcomes. We assessed the effects of mesenchymal proliferation in cancer gene expression data (Supplemental Figure S5). Using overall survival as an endpoint, and applying strict significance thresholds (see Methods), we found that three cancer types contained EMT-inflammation associated genes that predicted clear differences between patient groups. The three significant tumor types were bladder (BLCA), uterine (UCEC), and liver cancer (LIHC).

Next we modeled the effects of relevant gene sets on invasion-free survival, using the disease-free interval (DFI) as the endpoint. We clustered patients from each tumor type into two groups (high or low) based on their DFI (see Methods); these data contain 184 patients for BLCA, 114 patients for UCEC, and 311 patients for LIHC. For this clustering, the predictive accuracies (obtained by leave-one-out cross-validation) were 0.68 (BLCA), 0.69 (UCEC), and 0.621 (LIHC). We note that it is encouraging to obtain this level of accuracy on what is a challenging task: unsupervised prediction of survival differences, using a set of only 40 genes and with only ∼100 patients per tumor type. Several of the other 11 cancer types tested also displayed mesenchymal proliferation-associated effects, however these cancers were filtered out at the previous step, as they did not meet the significance thresholds set above.

We used Gaussian process classification to identify relationships between mesenchymal proliferation genes based on their ability to predict invasiveness (high or low DFI). We focus on interactions within the *TGF-β* and Wnt pathways, given their important roles in mediating EMT [56], and regulating cancer stem cell identity [57, 58]. *TGF-β* and Wnt pathways interact at multiple points, including through the *LEF1/TCF* complex [59], and via dimerization of their respective membrane-bound receptors [60]. The left-hand column of Figs. 5 and 6 shows the density of patient data, and thus the region (cyan box) to focus predictions on, over a two-dimensional gene expression region. The middle column shows the probability of high DFI over the same region, and the right-hand column depicts a slice through the co-expression plot.

**Figure 5:**
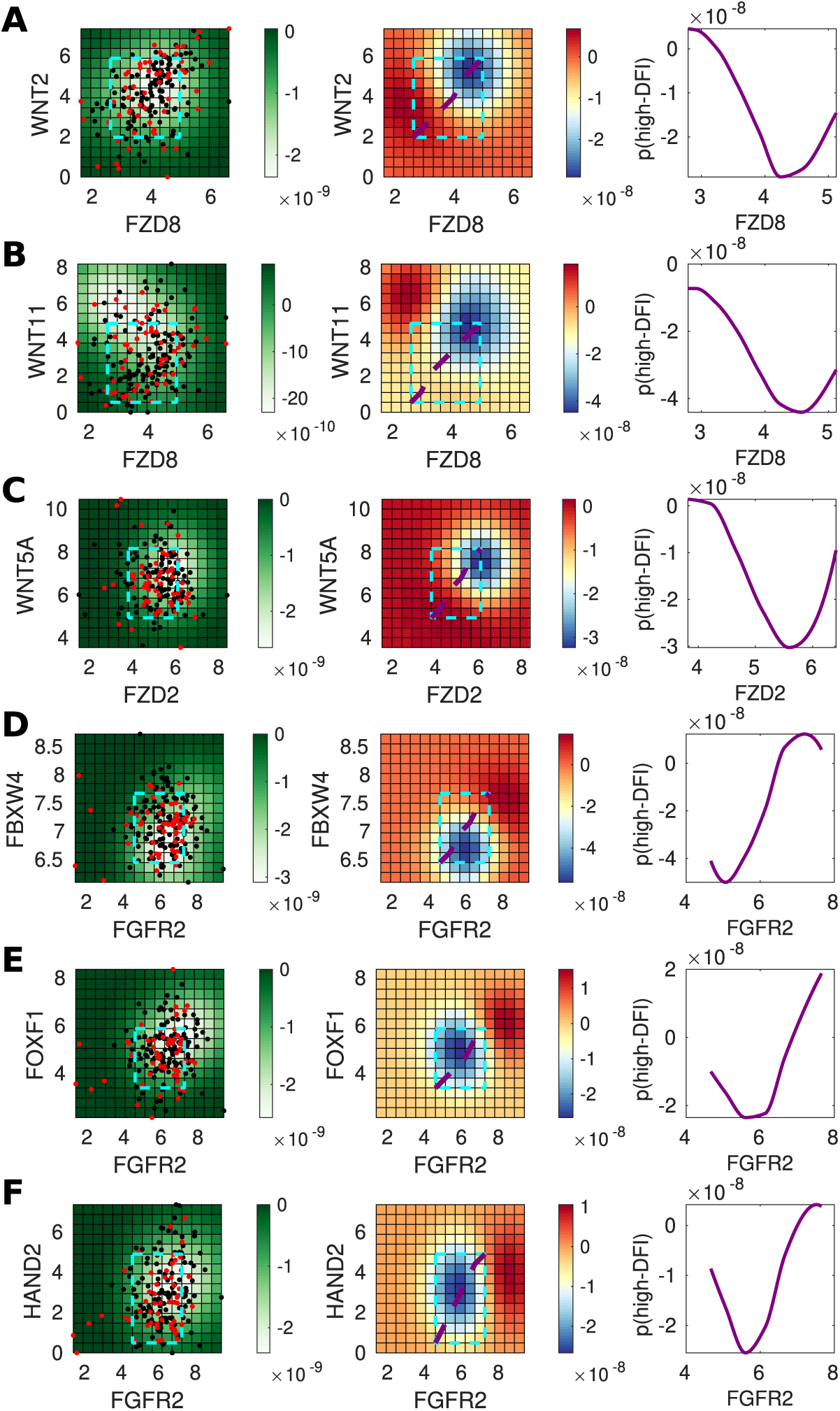
Genes predictive of invasiveness in BLCA. **A.** For gene pair *WNT2* and *FZD8*, the left panel shows the posterior variance on log-log expression plot of the predicted probability overlaid with patient samples (red = low-DFI, black = high-DFI), 90% confidence interval box drawn for standardized expression values (cyan); middle panel: posterior log probability of high-DFI over the same region as left, where the diagonal line (purple) shows the co-expression trend (diagonal line through the 90% CI of standardized expression values); right panel: posterior log probability of high-DFI plotted against expression of *FZD8*, values simulated along the diagonal (purple) corresponding to the middle panel. **B.** As above for *WNT11* and *FZD8*. **C.** As above for *WNT5A* and *FZD2*. **D.** As above for *FBXW4* and *FGFR2*. **E.** As above for *FOXF1* and *FGFR2*. **F.** As above for *HAND2* and *FGFR2*.

**Figure 6:**
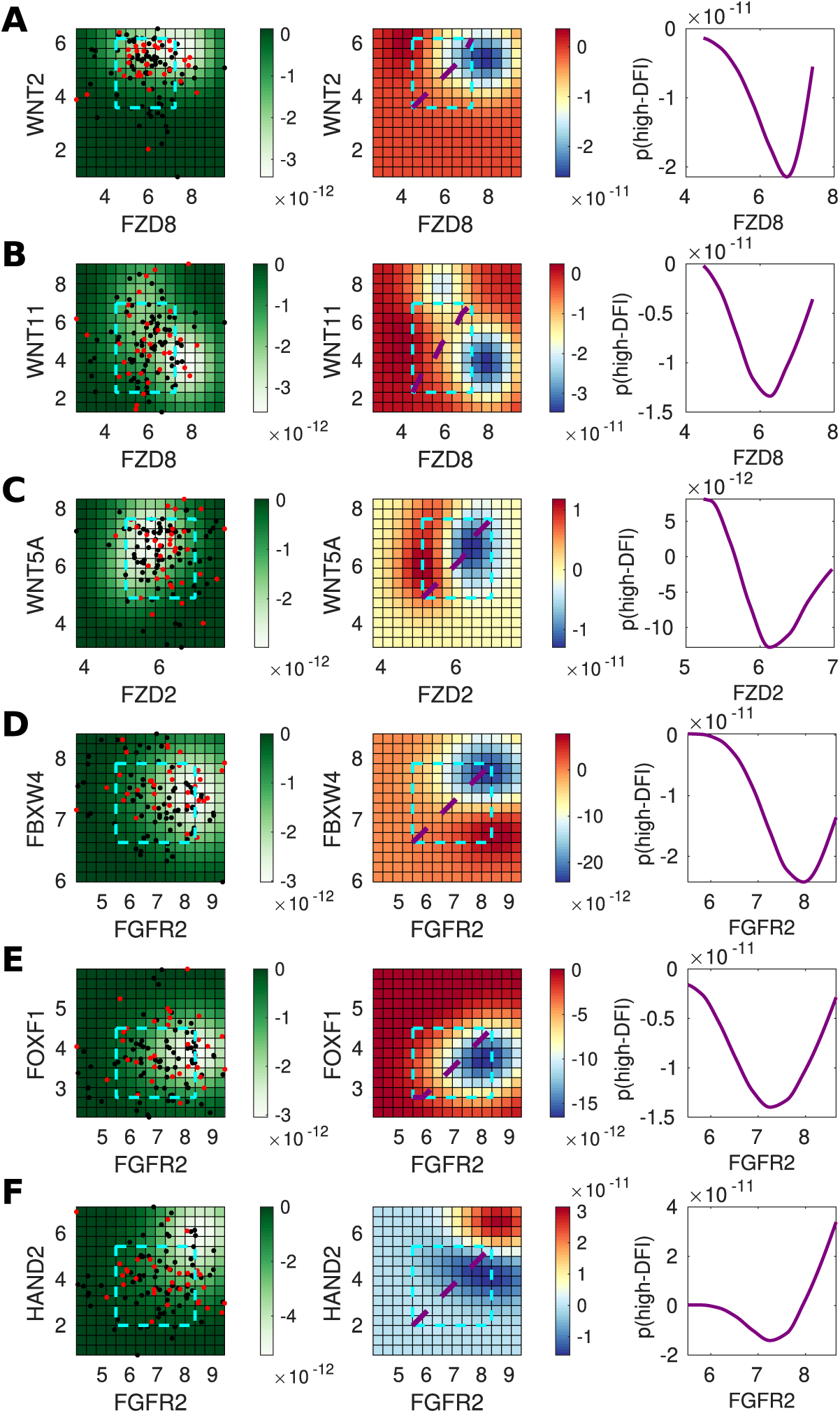
Genes predictive of invasiveness in UCEC. **A.** For gene pair *WNT2* and *FZD8*, theleft panel shows the posterior variance on log-log expression plot of the predicted probability overlaid with patient samples (red = low-DFI, black = high-DFI), 90% confidence interval box drawn for standardized expression values (cyan); middle panel: posterior log probability of high-DFI over the same region as left, where the diagonal line (purple) shows the co-expression trend (diagonal line through the 90% CI of standardized expression values); right panel: posterior log probability of high-DFI plotted against expression of *FZD8*, values simulated along the diagonal (purple) corresponding to the middle panel. **B.** As above for *WNT11* and *FZD8*. **C.** As above for *WNT5A* and *FZD2*. **D.** As above for *FBXW4* and *FGFR2*. **E.** As above for *FOXF1* and *FGFR2*. **F.** As above for *HAND2* and *FGFR2*.

In agreement with the literature, our results show that for both canonical and non-canonical Wnt signaling, higher levels of signaling leads to worse outcomes (Fig. 5A-C and Fig. 6A-C, summarized in the right-hand column): as the co-expression of the Wnt ligand and its receptor increases, the probability of a high DFI (better outcome) decreases. Studying the Wnt ligand-receptor predictions, though not monotonic in all cases, comparing low vs. high expression levels shows clear differences. This is the case for all three ligand-receptor pairs studied, for both BLCA and UCEC, with the exception in UCEC of *WNT11* and *FZD8* (Fig. 6B), where the co-expression effects are less clear. Overall, these predictions agree with expected tumorigenic roles for canonical [61, 62] and non-canonical [60, 63, 64] Wnt signaling in bladder (BLCA) and uterine (UCEC) cancers.

We identified several other gene pairs that can predict differences between high vs. low DFI patient groups (Fig. 5D-F and Fig. 6D-F). For the gene pairs (*FGFR2, FBXW4*) and (*FGFR2, FOXF1*) we see that a “goldilocks” region exists that most benefits survival - displaying striking similarity to the predicted mesenchymal effects in our model (Fig. 4E). We see that the predictions made by our modeling of single gene effects are in close agreement with the literature. Based on the agreement seen, we studied several new predictions made regarding the joint effects of co-expression on patient outcomes.

In Fig. 5D (middle panel) we show that a strong tumor suppressor effect of *FGFR2* in bladder cancer is predicted by our model. Although FGF signaling plays opposing roles in cancer, and FGFs can be up-regulated in tumors relying on FGF signaling for growth [65], *FGFR2* is implicated as a tumor suppressor in prostate and bladder cancer [66, 67]. We also predict a suppressive role for *FBXW4* (Fig. 5D, middle panel): for given *FGFR2* expression, increasing *FBXW4* leads to better outcomes. This agrees with literature suggesting that *FBXW4* is lost or mutated in almost 40% of urinary tract cancers [68]. Co-expression analysis also gives a new prediction: *FGFR2* and *FBXW4* act synergistically in BLCA, such that higher expression levels of both leads to greater outcomes than high expression of either gene alone (5D, middle & right panels). In comparison, for UCEC, the role is less clear, although the tumorigenic effect of *FGFR2* in uterine cancer is evident at high levels of *FBXW4* (6D), which is in line with previous studies reporting mutations that provide constitutive activation of *FGFR2* in a subset of endometrial cancer [69].

We highlight two further notable predictions. First, for BLCA, high *FGFR2* and *FOXF1* co-expression improves patient outcomes (Fig. 5E). The tumor suppressor *FOXF1* is a p53 target and it is epigenetically silenced in breast cancer [70, 71], however it was not previously known to have a tumor-suppressive role in BLCA either alone or co-expressed with *FGFR2*. This effect is not seen for UCEC (Fig. 6E), where our model predicts that the effects of *FGFR2* are tumorigenic in this region, but not affected by *FOXF1* expression (i.e. no significant differences between high and low co-expression). The second new prediction is that high co-expression of *FGFR2* and *HAND2* improves outcomes in UCEC (Fig. 6F); in contrast to the effects seen for the co-expression of *FGFR2* with either *FBXW4* or *FOXF1* (Fig. 6D-E), where, in each case, higher *FGFR2* expression led to worse outcomes. *HAND2* antagonizes FGF-dependent epithelial cell proliferation and is a critical regulatory component of both healthy and cancerous endometrial proliferation [72, 73]. For BLCA, we observe a less pronounced although still suppressive effect due to *HAND2* expression (Fig. 5F), in line with previous reports [74].

## 4 Discussion

Despite the importance of studying interactions between cancer and the immune system, as well as studying the effects of EMT on cancer, there has not previously, to the best of our knowledge, been a model developed that combines all three of these components. Here we studied cancer, the immune system, and EMT, during the progression from an in situ tumor to invasive disease. We saw this as a particularly pressing need given the shared factors influencing all these components, including *TGF-β* and Wnt signaling. We used an individual cell-based model framework to describe the multiscale processes leading to invasive disease, and we compared model predictions with a novel TCGA analysis framework to predict the effects mesenchymal phenotype-associated genes.

We found that the model recapitulated invasion-free survival dynamics. Using global parameter sensitivity analysis, we identified parameters exerting key control over model behavior. Focusing on these led us to identify that increasing mesenchymal immune evasion and increasing Treg *TGF-β* production both lead to shorter invasion-free survival times. However, varying the level of inflammation led to paradoxical effects with regards to mesenchymal growth arrest: under regimes with periods of low inflammation, an optimal level of mesenchymal growth arrest can improve outcomes and maximize the invasion-free survival. To capture the essential characteristics of the model, we summarized in silico patient studies with a single parameter: the invasion-free survival time. There are, of course, many trajectories that result in progression to invasion. Further analysis of the transient cell dynamics in tumors in situ and during progression is needed to gain greater insight into the EMT-associated dynamics of cancer.

We tested the model prediction that mesenchymal phenotypes play key roles in tumor invasiveness through a novel TCGA data analysis framework. In support of this prediction, we found that mesenchymal-associated genes controlled outcomes and predicted differences between high vs. low disease-free interval patient groups. This analysis yielded predictions of the effects of single genes or gene pairs, many of which corresponded to known effects, including the effects of both canonical and non-canonical Wnt signaling on tumor progression. Our modeling also predicted opposing roles for FGF signaling in bladder and uterine cancers: where *FGFR2* exerts a tumor-suppressor effect in bladder cancer yet a tumorigenic effect in uterine cancer. Evidence for these opposing roles already exists in the literature, but notably, through our modeling, we also predict entirely novel interactions between *FGFR2* and other transcription factors (*FBXW4, FOXF1*, and *HAND2*) that act to enhance or suppress the effects of *FGFR2* alone, and could offer significant novel therapeutic strategies.

In future work, further development of the inflammation module is important given the large and at times paradoxical roles that the inflammatory state exerts on tumor cells and invasion-free survival. Currently, inflammation is modeled as independently cycling between high and low schemes, yet several known factors contribute to the inflammatory state. For example, model extensions could assume that the level of inflammation depends on the number of and the degree of mutations that tumor cells harbor. The competing effects that *TGF-β* exerts on the tumor and its microenvironment also warrant further investigation. We found that – below a certain threshold – reduction of *TGF-β* increases the time to invasion, i.e. reducing *TGF-β* in the TME benefits survival. Experimental work in support of this result includes a study of *TGF-β* tumor suppression in pancreatic cancer through the promotion of EMT [75]. The *TGF-β* pathway is however implicated in numerous other cellular signaling processes besides EMT; changing *TGF-β* concentration even in a local environment could have large off-target effects. Indeed, it has been shown that *TGF-β* promotes invasion and heterogeneity while suppressing cell proliferation in squamous cell carcinoma [76]. To account for this complex signaling, future work should incorporate the effects of signaling factors downstream of *TGF-β* on the cancer dynamics. It is also important to note that in this model, EMT is initiated entirely by *TGF-β*. While *TGF-β* does play a large role in cancer EMT, it is by no means the only factor at play; in reality, cells must contend with and respond to a milieu of EMT-associated signals.

Tumor heterogeneity often helps the tumor to evade immune effects and complicates our approaches to treatment. Rigorous study of the consequences of the increased heterogeneity that follows disease incidence (i.e. decanalization [77]) is too-often sidelined, despite mounting evidence in support of its prominent role in cancer evolution [78–80]. Despite these challenges, great progress in predicting disease complexity continues to be made. As we are rapidly approaching a new generation of immunotherapies, it is these very complexities that we must better understand in order to control or eradicate the disease.

## Acknowledgements

Q.N. would like to acknowledge partial support for this work from National Institutes of Health grants R01GM123731, U01AR073159, and U54-CA217378; National Science Foundation grants DMS1562176 and DMS1763272; Simons Foundation grant (594598); and the Jayne Koskinas Ted Giovanis Foundation for Health and Policy joint with the Breast Cancer Research Foundation. A.L.M. would like to acknowledge partial support for this work from an American Cancer Society grant #IRG-16-181-57. M.K.K. was supported by the Mathematical, Computational and Systems Biology Predoctoral Training Grant T32 EB09418.

## Notes

### Competing Interest Statement

The authors have declared no competing interest.

