## Supplemental Methods and Tables for "Modeling the effects of EMT-immune dynamics on epithelial cancer progression"

### Supplementary Text and Tables for Modeling the competing effects of the immune system and EMT on epithelial cancers

**2 Definition of parameters specifying the model** **6**

**3 Parameter values used for simulation** **7**

**4 TCGA Analysis** **8**
4.1 Overall Survival . . . 9
4.1.1 Cox-PH Model . . . 9
4.2 Disease-Free Interval . . . 10

4.2.1 DFI is a Clinical Analogue of Tumor Time to Invasion 10
4.2.2 Ontology-Based Investigation of Proliferation Pathways 11
4.2.3 Binary Classification of DFI Endpoints . . . . . . . . 11

**5 Cox-PH Tables 14**
5.1 BLCA . . . . . . . . . . . . . . . . . . . . . . . . . . . . 14
5.2 CESC . . . . . . . . . . . . . . . . . . . . . . . . . . . . 14
5.3 COAD . . . . . . . . . . . . . . . . . . . . . . . . . . . . 14
5.4 ESCA . . . . . . . . . . . . . . . . . . . . . . . . . . . . 15
5.5 KIRP . . . . . . . . . . . . . . . . . . . . . . . . . . . . 15
5.6 LIHC . . . . . . . . . . . . . . . . . . . . . . . . . . . . 16
5.7 PAAD . . . . . . . . . . . . . . . . . . . . . . . . . . . . 17
5.8 UCEC . . . . . . . . . . . . . . . . . . . . . . . . . . . . 18

**6 KM Tables 20**

### 1 Model Description

#### 1.1 Tissue cell fate

During each cell cycle, every cell randomly is assigned a cell fate from the following options:

- proliferation
- apoptosis
- immune clearance (by NKs or CTLs)
- rest in $G_0$

For each cell, a weight is chosen for each option and these are normalized to probabilities which then are used to randomly determine what each cell does during the cell cycle.

##### 1.1.1 Proliferation

There are four factors that contribute to the weight of a cell to proliferate. The first is a base proliferation rate that all cells have, $p$. Second, if the cell has a mutation in the proliferation pathway ($\delta_P = 1$), then the weight for proliferation is proportionally increased by $\Delta_P$. Third, if the cell is mesenchymal ($\zeta = 1$), then the weight for proliferation is proportionally decreased by $\Delta_{\mathrm{MGA}}$, which stands for mesenchymal growth arrest. This lost proliferation for mesenchymal cells will later be used to increase their chance of resting. Fourth, there is a negative feedback of the cells on their own proliferation which is quantified by a Hill factor as a function of the tissue cell population, $N_C$, with EC50 term $K_0$. In total, the weight for proliferation is given by

$$\rho_P = p(1 + \delta_P \Delta_P)(1 - \zeta \Delta_{\text{MGA}}) \frac{K_0}{K_0 + N_C} \tag{2.1}$$

##### 1.1.2 Apoptosis

There are two factors that contribute to a cell's weight for undergoing apoptosis. There is a basal apoptosis rate that all cells experience, $d_C$ for death. Second, if the cell has a mutation in the apoptosis pathway ($\delta_A = 1$), then the weight for undergoing apoptosis is proportionally decreased by $\Delta_A$. In total, the weight for apoptosis is given by

$$\rho_A = d_C(1 - \delta_A \Delta_A) \tag{2.2}$$

##### 1.1.3 Immune Clearance

For both NK clearance and CTL clearance, the weights are built with the same factors but have different parameter values for NK and CTLs. First of all, the cell needs to be malignant ($\delta_{\text{MUT}} = 1$). Second, there is a Hill factor that captures the probability of an immune cell finding and interacting with the given tissue cell with EC50 term $K_1$. Third, NKs and CTLs have their own efficacy parameters, $E_{\text{NK}}$ and $E_{\text{CTL}}$, which can be understood as the rate of immune clearance given an immune cell has found the mutated cell. Fourth, there is a decreasing Hill factor based on the number of Treg cells present with EC50 term $K_2$. Finally, there are two factors that proportionally decrease the weight of immune clearance depending on if the cell has an immune evasion mutation ($\delta_{\text{IE}} = 1$) or if it is mesenchymal ($\zeta = 1$) with respective decreases $\Delta_{\text{IE}}$ and $\Delta_{\text{MIE}}$. In total, the weight of NK clearance is given by

$$\rho_{\text{NK}} = \delta_{\text{MUT}} \frac{N_{\text{NK}}}{N_C/K_1 + N_{\text{NK}}} \frac{E_{\text{NK}}}{1 + N_{\text{Treg}}/K_2} (1 - \delta_{\text{IE}} \Delta_{\text{IE}})(1 - \zeta \Delta_{\text{MIE}}) \tag{2.3}$$

A similar formula holds for CTLs with only the number of CTLs and their efficacy being different from the above equation.

##### 1.1.4 Rest in $G_0$

The weight associated with rest is taken as 1 except in the case of mesenchymal cells. Recall that mesenchymal cells had their proliferation rate decreased by $1 - \zeta \Delta_{\text{MGA}}$ (see Eq. 2.1). The biological assumption here is that mesenchymal cells instead of proliferating will instead rest, so this lost proliferation weight is added to the resting weight. Hence, the weight of rest is given by

$$\rho_R = 1 + \zeta p(1 + \delta_P \Delta_P)\Delta_{\text{MGA}} \frac{K_0}{K_0 + N_C} \tag{2.4}$$

Again, the reason for adding that term is due to the understanding that overall mesenchymal cells proliferate less as individual cells rest longer in the $G_0$ phase.

##### 1.1.5 Completing the Cell Cycle

After the cell fates are determined and the results reflected in the system, there are a few things that happen before the system moves on to a new cell cycle. First, the NK and CTL populations are reduced by the number of mutated cells they cleared. This represents the fact that individual immune cells lose efficacy as they carry out their effector functions. Second, all proliferating cells have a cell-specific probability of undergoing a driver mutation in one of the three pathways. If they do, one is randomly chosen among the three pathways and the pathway in that cell becomes altered. If the cell does not undergo a mutation, then its probability of mutation during subsequent cell cycles increases.

Finally, the EMT values for each cell is updated. This depends on the cells current EMT score and how much TGF-$\beta$ is currently in the system. The amount of TGF-$\beta$ absorbed by all cells is given by an increasing Hill function in terms of the TGF-$\beta$ in the TME. The saturation effect is to limit the amount of TGF-$\beta$ a cell can absorb in a given time interval. This quantity is then divided up randomly among the $N_C$ living cells via a normally distributed noise term to determine how much exogenous TGF-$\beta$ each cell receives during this cell cycle. Should this value, $\tau_i$ in Eq. 2.5, be negative, we interpret this as the cell losing TGF-$\beta$ to the TME and thus being more likely to undergo MET.

$$\tau_i = \frac{\tau_{\max}}{N_C} \frac{\tau/K_3}{1 + \tau/K_3} + X_i, \quad X_i \sim N(0, \sigma^2) \tag{2.5}$$

We then combine $\tau_i$ with the current EMT score of the cell, as a proxy for the endogenous TGF-$\beta$. Finally, if this quantity is large enough, the EMT score of the cell increases towards 1; otherwise, it decreases towards 0. Each cell then is relabeled as either epithelial or mesenchymal depending on its new EMT score and whether it is below or above the mesenchymal threshold. Thus, there are two main factors that determine if a cell will end a cell cycle as mesenchymal: concentration of TGF-$\beta$ in the system and the current EMT score of the cell.

Next, the amount of TGF-$\beta$ for the next cell cycle is determined by the number of mutated cells, $N_{\text{MUT}}$, and the number of Treg cells, $N_{\text{Treg}}$, each one producing a fixed amount of TGF-$\beta$. It is given by

$$\tau = \tau_{\text{MUT}} N_{\text{MUT}} + \tau_{\text{Treg}} N_{\text{Treg}} \tag{2.6}$$

Finally, the immune populations are updated. For the NKs, they obey the following differential equation:

$$N'_{\text{NK}} = \sigma_{\text{NK}} - d_{\text{NK}} N_{\text{NK}} \tag{2.7}$$

which is discretized to

$$N_{\text{NK}}(k+1) = \left( N_{\text{NK}}(k) - \frac{\sigma_{\text{NK}}}{d_{\text{NK}}} \right) \exp(-d_{\text{NK}} \Delta t) + \frac{\sigma_{\text{NK}}}{d_{\text{NK}}} \tag{2.8}$$

For CTLs and Tregs, they rely on malignant cells being cleared before they can be activated. Let $N^*_{\text{MUT}}(k)$ represent the number of malignant cells cleared by the immune system during cell cycle $k$. In addition, Treg recruitment is upregulated by TGF-$\beta$, which will be incorporated via a Hill function with EC50 term $K_4$. We choose the following differential equations to govern the CTL and Treg populations:

$$\begin{aligned} N'_{\text{CTL}} &= \sigma_{\text{CTL}} N^*_{\text{MUT}} - d_{\text{CTL}} N_{\text{CTL}} \\ N'_{\text{Treg}} &= \sigma_{\text{Treg}} N^*_{\text{MUT}} \frac{\tau}{1 + \tau/K_4} - d_{\text{Treg}} N_{\text{Treg}} \end{aligned} \tag{2.9}$$

Discretized, these are:

$$\begin{aligned} N_{\text{CTL}}(k+1) &= (N_{\text{CTL}}(k) - \sigma_{\text{CTL}} N^*_{\text{MUT}}(k)/d_{\text{CTL}}) \exp(-d_{\text{CTL}} \Delta t) + \sigma_{\text{CTL}} N^*_{\text{MUT}}(k)/d_{\text{CTL}} \\ N_{\text{Treg}}(k+1) &= \left( N_{\text{Treg}}(k) - \frac{\sigma_{\text{Treg}} N^*_{\text{MUT}}(k)}{d_{\text{Treg}}} \frac{\tau(k)}{1 + \tau(k)/K_4} \right) \exp(-d_{\text{Treg}} \Delta t) \\ &\quad + \frac{\sigma_{\text{Treg}} N^*_{\text{MUT}}(k)}{d_{\text{Treg}}} \frac{\tau(k)}{1 + \tau(k)/K_4} \end{aligned} \tag{2.10}$$

#### 2 Definition of parameters specifying the model

| Name | Description |
|---|---|
| $p$ | proliferation rate of tissue cells |
| $d_C$ | death rate of tissue cells |
| $\Delta_{\text{MIE}}$ | mesenchymal immune evasion |
| $\Delta_{\text{MGA}}$ | mesenchymal growth arrest |
| $\Delta_A$ | mutant cells decreased apoptosis |
| $\Delta_{\text{IE}}$ | mutant cells increased immune evasion |
| $\Delta_P$ | mutant cells increased proliferation |
| $K_0$ | EC50 term for negative feedback of tissue cells on own proliferation |
| $K_1$ | EC50 term for probability of NK cell finding mutant cell |
| $K_2$ | EC50 term for Treg inhibition of cytotoxic functions |
| $K_3$ | EC50 term for how much TGF-$\beta$ each cell has |
| $K_4$ | EC50 term for TGF-$\beta$ activation of Tregs |
| $E_{\text{NK}}$ | rate of NKs clearing mutants |
| $E_{\text{CTL}}$ | rate of CTLs clearing mutants |
| $\sigma_{\text{NK}}$ | NK source rate |
| $\sigma_{\text{CTL}}$ | CTL source rate per cleared mutant cell |
| $\sigma_{\text{Treg}}$ | Treg source rate per cleared mutant cell |
| $d_{\text{NK}}$ | NK death rate |
| $d_{\text{CTL}}$ | CTL death rate |
| $d_{\text{Treg}}$ | Treg death rate |
| $k_{\text{EMT}}$ | EMT/MET rate |
| $\sigma$ | standard deviation of noise in TGF-$\beta$ each cell receives |
| $\tau_{\max}$ | max amount of TGF-$\beta$ any cell can receive |
| $\tau_{\text{MUT}}$ | rate of TGF-$\beta$ production by mutant cells |
| $\tau_{\text{Treg}}$ | rate of TGF-$\beta$ production by Treg |

Table S1: ]
The model parameter names and descriptions. Note that many of these values are affected by the inflammation state of the system.

#### 3 Parameter values used for simulation

| Name | Description | INFL Low Value | INFL High Value |
|---|---|---|---|
| $p$ | weight of proliferation for tissue cells | 0.28 | |
| $d_C$ | weight of apoptosis for tissue cells | 0.14 | |
| $\Delta_{\mathrm{MIE}}$ | MIE | 0.6 | |
| $\Delta_{\mathrm{MGA}}$ | MGA | 0.2 | |
| $\Delta_A$ | proportional decrease to weight of apoptosis for cells with mutated apoptosis pathway | 0.3 | |
| $\Delta_{\mathrm{IE}}$ | proportional increase to weight of immune evasion for cells with mutated immune evasion pathway | 0.48 | |
| $\Delta_P$ | proportional increase to weight of proliferation for cells with mutated proliferation pathway | 0.36 | |
| $K_0$ | EC50 term for negative feedback of tissue cells on own proliferation | 80 cells | |
| $K_1$ | EC50 term for probability of NK cell finding mutant cell | 8 cells | |
| $K_2$ | EC50 term for Treg inhibition of cytotoxic functions | 5 cells / volume | 0.025 cells / volume |
| $K_3$ | EC50 term for cumulative absorption of TGF-$\beta$ | 200 amount / volume | |
| $K_4$ | EC50 term for TGF-$\beta$ activation of Tregs | 50 amount / volume | |
| $E_{\mathrm{NK}}$ | weight of NKs clearing mutants | 10 | 30 |
| $E_{\mathrm{CTL}}$ | weight of CTLs clearing mutants | 200 | 600 |
| $\sigma_{\mathrm{NK}}$ | NK source rate | 1.3 cells / cycle | |
| $\sigma_{\mathrm{CTL}}$ | CTL source rate per cleared mutant cell | 100 cells / (cleared mutants $\times$ cycles) | |

| | | |
|---|---|---|
| $\sigma_{\text{Treg}}$ | Treg source rate per cleared mutant cell | 200 cells / (cleared mutants $\times$ concentration of TGF-$\beta$ $\times$ cycles) |
| $d_{\text{NK}}$ | NK death rate | 0.13 / cycle |
| $d_{\text{CTL}}$ | CTL death rate | 0.0260 / cycle |
| $d_{\text{Treg}}$ | Treg death rate | 0.0260 / cycle |
| $k_{\text{EMT}}$ | EMT/MET rate | 0.01 / concentration of TGF-$\beta$ |
| $\sigma$ | standard deviation of noise in TGF-$\beta$ each cell receives | 6 concentration of TGF-$\beta$ |
| $\tau_{\text{max}}$ | max amount of TGF-$\beta$ any cell can receive | 500 concentration of TGF-$\beta$ |
| $\tau_{\text{MUT}}$ | rate of TGF-$\beta$ production by mutant cells | 0.05 concentration of TGF-$\beta$ / cell / cycle |
| $\tau_{\text{Treg}}$ | rate of TGF-$\beta$ production by Treg | 0.5 concentration of TGF-$\beta$ / cell / cycle |
| | RP Cancer Line | 0.5 |
| | INFL High Duration | 30 cycles |
| | INFL Low Duration | 30 cycles |
| | Mes Threshold | 0.7 |
| | maximum initial mutation damage after warmup | 0.01 |
| | increase in probability to mutate for non-mutating proliferating cells | 0.0001 |

Table S2: The model parameter names, descriptions, and values during both low and high inflammation. Parameters with only one value do not change with the inflammatory state.

### 4 TCGA Analysis

The Cancer Genome Atlas (TCGA) [7] provides multiple clinical endpoints, including overall survival (OS) and disease-free interval (DFI) [21]. In order to investigate the link between EMT, inflammation and invasive phenotype, we corroborate our model with clinical data from TCGA in a two pronged approach:

- **Overall Survival:** Test whether EMT and inflammation can jointly separate clinical cohorts based on the OS endpoint for a selection of cancer sub-types (Section 4.1).

- **Disease-Free Interval:** Identify pathway genes which regulate the proliferation/tumor-invasiveness axis in the context of a synergistic EMT/Inflam effect (Section 4.2).

Our analysis of tumor invasiveness due to mesenchymal growth arrest, in the context of EMT and inflammation takes place in two steps (fig S5.A-D): First, identify relevant cancers by A) defining sets of genes which represent some union of inflammation and EMT pathways while simultaneously having a quantitatively greater effect on overall survival than the component pathways acting alone, and B) simulating the dosage effect of proliferation markers on tumor invasiveness, for cancer types where a synergistic EMT/Inflam pathway was identified in (A). Following guidelines published in [21], we investigated the 14 TCGA tumor types recommended for both OS and DFI analysis.

#### 4.1 Overall Survival

##### 4.1.1 Cox-PH Model

While immunological interactions and EMT are known to be related [13], there is uncertainty regarding both the individual pathways which govern this dependence and the extent to which the interaction between inflammation and EMT is synergistic. Our approach identifies pathways by gene set (among all pairwise combinations of EMT and inflammation gene sets available from MSigDB [19]) for which the synergistic relationship between EMT and inflammation has a greater effect on OS than either process individually. For each combination of gene sets, we created three (one-dimensional) UMAP projections [23] of the data, one each from A) the EMT genes, B) the Inflammation genes, and C) the concatenation of (A) and (B). This yielded a three-dimensional projection of the data, on which we build a Cox proportional hazard model (CoxPH). This approach resembles a PCA-based approach introduced in [33] for SNP-based predictions. We identified several gene set combinations (combos) for which the global statistical significance (by likelihood ratio test) of the corresponding model was high ($p_{LR} \leq 5e-2$), as were all three predictors, but for which the hazard ratios for the concatenation embedding were at least 5% greater in magnitude than either EMT or inflammation alone. Prior to CoxPH analysis, the proportional hazard assumption was tested and only tumor-type/combos were retained whose Schoenfeld residual was equal to 0 [15]. Our screen identified 13 tumor-type/combos across 8 tumor types: urothelial bladder carcinoma (BLCA, table S3), Cervical Squamous Cell Carcinoma and Endocervical Adenocarcinoma (CESC, table S4), colon adenocarcinoma (COAD, S5), esophageal carcinoma (ESCA, S6), cervical kidney renal papillary cell carcinoma (KIRP, tables S7, S8), liver hepatocellular carcinoma (LIHC, tables S9,S10), pancreatic adenocarcinoma (PAAD, tables S11,S12), and uterine corpus endometrial carcinoma (UCEC, tables S13,S14,S15). See supplementary file "OS_test_results.csv" for full results (all tumor-type/combos).

In order to provide further confirmation of a relationship between survival in these four cancers and the synergistic activation of relevant pathways, we tested the separation (adjusted $p_{\text{log rank}} \leq 0.05$, [15]) of KM models fitted to subgroups defined unsupervised hierarchical density-based clustering [5, 11] (DBSCAN) of the UMAP-embedded combined gene set. We guided the unsupervised clustering by scaling down the minimum neighborhood size (starting with 30 patients) until the number of clusters was at least two. In addition to assigning cluster labels, DBSCAN determines outliers based on the the neighborhood structure of the graph [5,11]. In our KM models and in subsequent analysis of DFI prediction, these outliers were discarded in order to ensure that groups of patients were maximally homogeneous with respect to EMT/inflammation. A single gene set combo met these criteria for the following cancers: BLCA (table S16, survival plot fig.S6), LIHC (table S17, survival plot fig.S7), and UCEC (table S18, survival plot fig.S8).

This analysis robustly identified BLCA, LIHC, and UCEC as cancers for which synergistic interaction between EMT and inflammation is the primary driver of patient survival. Our approach has several advantages. First, we utilized the MSigDB resource [19] in order to optimize the search space over relevant pathways. This allows the large volume of prior knowledge encoded in this database to guide exploratory data analysis that would otherwise be impossible or impractical at the transcriptomic scale [33]. Second, our use of dimensionality-reduction provides the following two-fold advantage: clear interpretation of the synergistic response between EMT and INFLAM and the compression of the parameter space to only three predictors, which means that sensitive prediction of survival can be carried out on the limited number of primary tumor samples in our data set. In the sequel, we address the role of mesenchymal proliferation pathways in the invasiveness of these tumors, by utilizing the disease-free interval (DFI) endpoint [21], rather than the OS endpoint.

#### 4.2 Disease-Free Interval

##### 4.2.1 DFI is a Clinical Analogue of Tumor Time to Invasion

Our agent-based simulations cover the incremental progression from in-situ to invasive disease from a homogeneous initial point, whereas the data in TCGA address how cancer may progress following treatment, thus comparisons between model and data should be made carefully. Nonetheless recent clinical and experimental evidence suggests that core cellular tumor dynamics are at play both during the tumor progression addressed by the model, and post-treatment progression described in data from TCGA. Of particular note, the plasticity of tumor cells allows them to evade treatment by undergoing post-treatment processes resembling the de-novo appearance of cancer [29].

##### 4.2.2 Ontology-Based Investigation of Proliferation Pathways

As stated above, we predict that for certain EMT and inflammatory environments, the time to invasion is maximized by a specific proliferative regime, where the proliferative potential of a transformed tumor cell is being held in check by mesenchymal growth arrest programs. Therefore we investigate the timing of invasion as a function of proliferation by searching for proliferative regimes where the Disease Free Interval (DFI) is maximized for patients in remission after treatment. In contrast to the search-based strategy above, we used the Gene Ontology (GO) resource [1, 8] to select an appropriate pathway for this analysis. GO is designed to provide a semantic index of genes, allowing gene lists to be retrieved interactively by simply browsing its hierarchy. We selected GO:0010463 (Mesenchymal Cell Proliferation) for our analysis of proliferation-dependent DFI.

##### 4.2.3 Binary Classification of DFI Endpoints

Binarization of survival endpoints has previously been explored [3, 6, 9, 14, 16, 18, 20]. In contrast to previous approaches which utilize a pre-determined time threshold for the response (e.g. early and late relapse), we utilized an imputed high/low risk classification scheme based on a two-component Gaussian mixture model, which implicitly deals with cancer-specific thresholds. This approach was motivated by the observation that in all three cases, the DFI exibited multiple modes with cancer-specific thresholds: BLCA (fig.S9), LIHC (fig.S11), UCEC (fig.S10). Under this scheme, a tumor with a short DFI represents highly invasive disease for which the time-to-invasion is short.

**Summary of the Model**

The log counts from TCGA bulk mRNA sequencing for 52 genes are used to predict the computed DFI-class for each patient. The list of genes includes the 42 human genes from GO:0010463 "mesenchymal cell proliferation" (which omits all but LRP5 among known receptors for WNT2/11/5A) augmented with missing receptors for those Wnts: FZD2, FZD4, FZD6, FZD8, ROR1, ROR2, RYK, and LRP6. The model encodes the response $y$ as either $+1$ or -1 (Eq. 1) for high-DFI and low-DFI respectively and is fitted via a generalized form of Bayesian logistic regression [27] using the expression levels of these genes as predictors. Since the likelihood (Eq. 1) is non-Gaussian, the posterior (Eq. 3) becomes analytically intractible, so expectation propagation (EP) is used to approximate it during inference and hyperparameter optimization [27]. Inference and hyperparameter optimization were performed using the gpstuff toolbox for MATLAB [30].

Our model simultaneously considers the (gene-expression of) multiple distinct biological pathways using the product of squared-exponential kernels over the predictor genes (Eq. 4). Within the context of the GP classifier, this kernel specifies the covariance of the joint distribution over any subset of the input data. The constant term $\sigma_0$, magnitude $\sigma^2$, and gene-wise length scale $\lambda_d$ are given priors (Eq. 5, 6, 7, 8) which facilitate the discovery of their MAP values by EP. This basic structure assumes little prior knowledge about the predictors, while offering good out of sample prediction accuracy.

$$p(\boldsymbol{y}|\boldsymbol{f}) = \prod_{i=1}^{n} \frac{1}{1 + exp(y_i f_i)} \tag{1}$$

$$p(\boldsymbol{f}|\boldsymbol{X}, \theta) = \mathcal{N}(\boldsymbol{f}|\boldsymbol{0}, \boldsymbol{K}) \tag{2}$$

$$p(\boldsymbol{f}|\boldsymbol{X}, \boldsymbol{y}, \theta) = \frac{p(\boldsymbol{f}|\boldsymbol{X}, \theta)}{p(\boldsymbol{X}|\theta)} \prod_{i=1}^{n} p(y_i|f_i) \tag{3}$$

$$\boldsymbol{K}(\boldsymbol{x}, \boldsymbol{x'}) = \sigma_0 + \sigma^2 exp\left[-\frac{1}{2}\sum_{d=1}^{D}\left(\frac{\boldsymbol{x} - \boldsymbol{x'}}{\lambda_d}\right)^2\right] \tag{4}$$

$\boldsymbol{x}$ and $\boldsymbol{x'}$ any two patients

$$log(\sigma_0) \sim \mathcal{N}(0, 0.1) \tag{5}$$

$$log(\sigma^2) \sim \mathcal{N}(1, 0.25) \tag{6}$$

$$log(\lambda_d) \sim \mathcal{N}(1, \Sigma_0) \tag{7}$$

$$\Sigma_0 \sim \mathcal{IG}(3, 1) \tag{8}$$

This model achieves very high ($\sim 1$) LOO-CV accuracy on the training data, so it is instructive to measure its performance relative to linear SVM on the sub-cohorts (noisy resamplings of the patient data) used for clustering. the average classification performance over all 1000 subcohorts is shown in the following table:

| Type | $-log(p(y))$ | Naive | Linear SVM | GP |
|---|---|---|---|---|
| LIHC | 197.800 | 0.582 | 0.582 | 1.0 |
| BLCA | 106.053 | 0.686 | 0.689 | 1.0 |
| UCEC | 62.321 | 0.708 | 0.726 | 1.0 |

Above, we list the leave-one-out cross-validated (LOO-CV) classification accuracy in each case for GP, and the 5-fold cross-validation accuracy for linear SVM, computed using the MATLAB Optimization Toolbox. The LOO-CV approach of [26] utilizing the cavity distribution of the EP likelihood approximation is utilized for tractability. This approach aims to discover the out of sample prediction accuracy for the model while simultaneously using all the data [31]. Compared to linear SVM [12], the GP classifier for all cancers achieves 100% LOO-CV, while SVM achieves only a modest improvement over naive (selecting high-DFI for all patients) for BLCA and UCEC, while failing improve the naive estimate for LIHC. This latter result is consistent with the higher negative log marginal likelihood $(-log(p(y)))$ for LIHC, indicating that the association between our chosen markers and the DFI endpoint is less justified. Therefore, LIHC was excluded from further analysis.

### 5 Cox-PH Tables

#### 5.1 BLCA

| | *Dependent variable:* |
|---|---|
| | time |
| EMT | −0.045** |
| | (0.018) |
| INFLAM | 0.025*** |
| | (0.009) |
| BOTH | 0.082*** |
| | (0.021) |
| Observations | 401 |
| $R^2$ | 0.056 |
| Max. Possible $R^2$ | 0.991 |
| Log Likelihood | −924.075 |
| Wald Test | 22.290*** (df = 3) |
| LR Test | 23.300*** (df = 3) |
| Score (Logrank) Test | 22.401*** (df = 3) |
| *Note:* | *p<0.1; **p<0.05; ***p<0.01 |

Table S3: Gotzman EMT vs. GO Pos Acute Inflam Ant
GOTZMANN_EPITHELIAL_TO_MESENCHYMAL_TRANSITION_UP
vs.
GO_POSITIVE_REGULATION_OF_ACUTE_INFLAMMATORY_RESPONSE_TO_ANTIGENIC_STIMULUS

#### 5.2 CESC

| | *Dependent variable:* |
|---|---|
| | time |
| EMT | −0.100*** |
| | (0.032) |
| INFLAM | −0.067*** |
| | (0.023) |
| BOTH | 0.092** |
| | (0.041) |
| Observations | 291 |
| $R^2$ | 0.060 |
| Max. Possible $R^2$ | 0.910 |
| Log Likelihood | −340.512 |
| Wald Test | 18.320*** (df = 3) |
| LR Test | 18.083*** (df = 3) |
| Score (Logrank) Test | 18.630*** (df = 3) |
| *Note:* | *p<0.1; **p<0.05; ***p<0.01 |

Table S4: GO Pos EMT vs. GO Leuk Act
GO_POSITIVE_REGULATION_OF_EPITHELIAL_TO_MESENCHYMAL_TRANSITION
vs.
GO_LEUKOCYTE_ACTIVATION_INVOLVED_IN_INFLAMMATORY_RESPONSE

#### 5.3 COAD

| | *Dependent variable:* |
|---|---|
| | time |
| EMT | $-0.142^{***}$ |
| | (0.041) |
| INFLAM | $0.067^{**}$ |
| | (0.026) |
| BOTH | $0.126^{***}$ |
| | (0.048) |
| Observations | 276 |
| $R^2$ | 0.048 |
| Max. Possible $R^2$ | 0.904 |
| Log Likelihood | −316.811 |
| Wald Test | $15.000^{***}$ (df = 3) |
| LR Test | $13.455^{***}$ (df = 3) |
| Score (Logrank) Test | $14.372^{***}$ (df = 3) |
| *Note:* | $^{*}$p<0.1; $^{**}$p<0.05; $^{***}$p<0.01 |

Table S5: Hollern EMT Breast vs. GO Leuk Act
HOLLERN_EMT_BREAST_TUMOR_UP
vs.
GO_LEUKOCYTE_ACTIVATION_INVOLVED_IN_INFLAMMATORY_RESPONSE

#### 5.4 ESCA

| | *Dependent variable:* |
|---|---|
| | time |
| EMT | $0.086^{**}$ |
| | (0.038) |
| INFLAM | $-0.113^{***}$ |
| | (0.042) |
| BOTH | $0.185^{***}$ |
| | (0.064) |
| Observations | 183 |
| $R^2$ | 0.052 |
| Max. Possible $R^2$ | 0.972 |
| Log Likelihood | −321.566 |
| Wald Test | $9.760^{**}$ (df = 3) |
| LR Test | $9.838^{**}$ (df = 3) |
| Score (Logrank) Test | $9.802^{**}$ (df = 3) |
| *Note:* | $^{*}$p<0.1; $^{**}$p<0.05; $^{***}$p<0.01 |

Table S6: GO Cardiac EMT vs. GO Neg Acute Inf

GO_CARDIAC_EPITHELIAL_TO_MESENCHYMAL_TRANSITION
vs.
GO_NEGATIVE_REGULATION_OF_ACUTE_INFLAMMATORY_RESPONSE

#### 5.5 KIRP

| | *Dependent variable:* |
|---|---|
| | time |
| EMT | $-0.090^{**}$ |
| | (0.041) |
| INFLAM | $-0.040^{***}$ |
| | (0.015) |
| BOTH | $0.103^{**}$ |
| | (0.040) |
| Observations | 287 |
| $R^2$ | 0.101 |
| Max. Possible $R^2$ | 0.769 |
| Log Likelihood | −194.964 |
| Wald Test | $35.480^{***}$ (df = 3) |
| LR Test | $30.644^{***}$ (df = 3) |
| Score (Logrank) Test | $41.090^{***}$ (df = 3) |
| *Note:* | $^{*}$p<0.1; $^{**}$p<0.05; $^{***}$p<0.01 |

Table S7: GO Reg EMT vs. GO Neg Acute Inf

GO_REGULATION_OF_EPITHELIAL_TO_MESENCHYMAL_TRANSITION
vs.
GO_NEGATIVE_REGULATION_OF_ACUTE_INFLAMMATORY_RESPONSE

| | *Dependent variable:* |
|---|---|
| | time |
| EMT | $-0.129^{***}$ |
| | (0.033) |
| INFLAM | $-0.068^{***}$ |
| | (0.026) |
| BOTH | $0.121^{***}$ |
| | (0.030) |
| Observations | 287 |
| $R^2$ | 0.113 |
| Max. Possible $R^2$ | 0.769 |
| Log Likelihood | −193.046 |
| Wald Test | $44.270^{***}$ (df = 3) |
| LR Test | $34.480^{***}$ (df = 3) |
| Score (Logrank) Test | $52.588^{***}$ (df = 3) |
| *Note:* | $^{*}$p<0.1; $^{**}$p<0.05; $^{***}$p<0.01 |

Table S8: GO Reg EMT vs. GO Neg Inf

GO_REGULATION_OF_EPITHELIAL_TO_MESENCHYMAL_TRANSITION
vs.
GO_NEGATIVE_REGULATION_OF_INFLAMMATORY_RESPONSE

#### 5.6 LIHC

| | *Dependent variable:* |
|---|---|
| | time |
| EMT | −0.029** |
| | (0.012) |
| INFLAM | −0.159** |
| | (0.065) |
| BOTH | 0.181*** |
| | (0.062) |
| Observations | 364 |
| $R^2$ | 0.058 |
| Max. Possible $R^2$ | 0.974 |
| Log Likelihood | −654.198 |
| Wald Test | 20.970*** (df = 3) |
| LR Test | 21.748*** (df = 3) |
| Score (Logrank) Test | 21.321*** (df = 3) |
| *Note:* | *p<0.1; **p<0.05; ***p<0.01 |

Table S9: GO Cardiac EMT vs. Zhou Inf FIMA Up

GO_REGULATION_OF_CARDIAC_EPITHELIAL_TO_MESENCHYMAL_TRANSITION
vs.
ZHOU_INFLAMMATORY_RESPONSE_FIMA_UP

| | *Dependent variable:* |
|---|---|
| | time |
| EMT | −0.026** |
| | (0.012) |
| INFLAM | −0.027*** |
| | (0.010) |
| BOTH | 0.033** |
| | (0.017) |
| Observations | 364 |
| $R^2$ | 0.046 |
| Max. Possible $R^2$ | 0.974 |
| Log Likelihood | −656.570 |
| Wald Test | 16.730*** (df = 3) |
| LR Test | 17.005*** (df = 3) |
| Score (Logrank) Test | 17.070*** (df = 3) |
| *Note:* | *p<0.1; **p<0.05; ***p<0.01 |

Table S10: GO Reg EMT Endo vs. GO Mac Inf Prot 1 Alpha

GO_REGULATION_OF_EPITHELIAL_TO_MESENCHYMAL_TRANSITION_INVOLVED_IN_ENDOCARDIAL_CUSHION_FORMATION
vs.
GO_MACROPHAGE_INFLAMMATORY_PROTEIN_1_ALPHA_PRODUCTION

#### 5.7 PAAD

| | *Dependent variable:* |
|---|---|
| | time |
| EMT | −0.036** |
| | (0.018) |
| INFLAM | −0.063** |
| | (0.029) |
| BOTH | 0.052*** |
| | (0.018) |
| Observations | 177 |
| $R^2$ | 0.061 |
| Max. Possible $R^2$ | 0.991 |
| Log Likelihood | −407.162 |
| Wald Test | 10.610** (df = 3) |
| LR Test | 11.187** (df = 3) |
| Score (Logrank) Test | 10.742** (df = 3) |
| *Note:* | *p<0.1; **p<0.05; ***p<0.01 |

Table S11: Alonso Met EMT Down vs. GO Neg Reg Inf
ALONSO_METASTASIS_EMT_DN
vs.
GO_NEGATIVE_REGULATION_OF_INFLAMMATORY_RESPONSE

| | *Dependent variable:* |
|---|---|
| | time |
| EMT | −0.099*** |
| | (0.033) |
| INFLAM | −0.043** |
| | (0.022) |
| BOTH | 0.070** |
| | (0.029) |
| Observations | 177 |
| $R^2$ | 0.099 |
| Max. Possible $R^2$ | 0.991 |
| Log Likelihood | −403.487 |
| Wald Test | 17.620*** (df = 3) |
| LR Test | 18.536*** (df = 3) |
| Score (Logrank) Test | 17.776*** (df = 3) |
| *Note:* | *p<0.1; **p<0.05; ***p<0.01 |

Table S12: Jechlinger EMT Down vs. GO Pos Reg Cyto Prod
JECHLINGER_EPITHELIAL_TO_MESENCHYMAL_TRANSITION_DN
vs.
GO_POSITIVE_REGULATION_OF_CYTOKINE_PRODUCTION_INVOLVED_IN_INFLAMMATORY_RESPONSE

#### 5.8 UCEC

| | *Dependent variable:* |
|---|---|
| | time |
| EMT | $-0.158^{***}$ |
| | (0.045) |
| INFLAM | $-0.140^{**}$ |
| | (0.062) |
| BOTH | $0.216^{***}$ |
| | (0.075) |
| Observations | 174 |
| $R^2$ | 0.081 |
| Max. Possible $R^2$ | 0.789 |
| Log Likelihood | −127.984 |
| Wald Test | $14.660^{***}$ (df = 3) |
| LR Test | $14.652^{***}$ (df = 3) |
| Score (Logrank) Test | $14.971^{***}$ (df = 3) |
| *Note:* | $^{*}$p<0.1; $^{**}$p<0.05; $^{***}$p<0.01 |

Table S13: Alonso Met EMT Up vs. Fulcher Inf Resp Lectin LPS Down
ALONSO_METASTASIS_EMT_UP
vs.
FULCHER_INFLAMMATORY_RESPONSE_LECTIN_VS_LPS_DN

| | *Dependent variable:* |
|---|---|
| | time |
| EMT | $-0.179^{***}$ |
| | (0.054) |
| INFLAM | $-0.047^{**}$ |
| | (0.019) |
| BOTH | $0.232^{***}$ |
| | (0.079) |
| Observations | 174 |
| $R^2$ | 0.088 |
| Max. Possible $R^2$ | 0.789 |
| Log Likelihood | −127.269 |
| Wald Test | $11.950^{***}$ (df = 3) |
| LR Test | $16.082^{***}$ (df = 3) |
| Score (Logrank) Test | $12.639^{***}$ (df = 3) |
| *Note:* | $^{*}$p<0.1; $^{**}$p<0.05; $^{***}$p<0.01 |

Table S14: GO Cardiac EMT vs. GO Reg Inf Resp Wound
GO_CARDIAC_EPITHELIAL_TO_MESENCHYMAL_TRANSITION
vs.
GO_REGULATION_OF_INFLAMMATORY_RESPONSE_TO_WOUNDING

| | *Dependent variable:* |
|---|---|
| | time |
| EMT | $-0.038^{**}$ |
| | (0.019) |
| INFLAM | $-0.172^{***}$ |
| | (0.044) |
| BOTH | $0.174^{***}$ |
| | (0.050) |
| Observations | 174 |
| $R^2$ | 0.094 |
| Max. Possible $R^2$ | 0.789 |
| Log Likelihood | $-126.768$ |
| Wald Test | $21.670^{***}$ (df = 3) |
| LR Test | $17.084^{***}$ (df = 3) |
| Score (Logrank) Test | $16.551^{***}$ (df = 3) |
| *Note:* | $^{*}$p<0.1; $^{**}$p<0.05; $^{***}$p<0.01 |

Table S15: GO Card EMT vs. Wunder Inf Resp Chol Up
GO_CARDIAC_EPITHELIAL_TO_MESENCHYMAL_TRANSITION
vs.
WUNDER_INFLAMMATORY_RESPONSE_AND_CHOLESTEROL_UP

### 6 KM Tables

| | N | Observed | Expected | (O-E)^2/E | (O-E)^2/V |
|---|---|---|---|---|---|
| Cluster 1 | 118 | 35.00 | 54.28 | 6.85 | 11.61 |
| Cluster 2 | 52 | 25.00 | 19.29 | 1.69 | 1.99 |
| Cluster 3 | 38 | 20.00 | 15.50 | 1.31 | 1.49 |
| Cluster 4 | 93 | 54.00 | 44.93 | 1.83 | 2.79 |

p=0.008

Table S16: Gotzman EMT vs. GO Pos Acute Inflam Ant
GOTZMANN_EPITHELIAL_TO_MESENCHYMAL_TRANSITION_UP
vs.
GO_POSITIVE_REGULATION_OF_ACUTE_INFLAMMATORY_RESPONSE_TO_ANTIGENIC_STIMULUS

| | N | Observed | Expected | (O-E)^2/E | (O-E)^2/V |
|---|---|---|---|---|---|
| Cluster 1 | 160 | 69.00 | 49.64 | 7.55 | 15.43 |
| Cluster 2 | 110 | 32.00 | 51.36 | 7.30 | 15.43 |

p=5e-8

Table S17: GO Reg EMT Endo vs. GO Mac Inf Prot 1 Alpha
GO_REGULATION_OF_EPITHELIAL_TO_MESENCHYMAL_TRANSITION_INVOLVED_IN_ENDOCARDIAL_CUSHION_FORMATION
vs.
GO_MACROPHAGE_INFLAMMATORY_PROTEIN_1_ALPHA_PRODUCTION

| | N | Observed | Expected | (O-E)^2/E | (O-E)^2/V |
|---|---|---|---|---|---|
| Cluster 1 | 88 | 21.00 | 14.86 | 2.54 | 5.12 |
| Cluster 2 | 76 | 9.00 | 15.14 | 2.49 | 5.12 |

p=0.02

Table S18: GO Cardiac EMT vs. GO Reg Inf Resp Wound
GO_CARDIAC_EPITHELIAL_TO_MESENCHYMAL_TRANSITION
vs.
GO_REGULATION_OF_INFLAMMATORY_RESPONSE_TO_WOUNDING

| Gene(s) | Cancer | Effect | Eff.Type | Citation |
|---|---|---|---|---|
| Wnts | Both | Onc | NA | [2, 4, 17, 24, 25] |
| FGFR2 | BLCA | Supp | NA | [28] |
| FGFR2 | UCEC | Onc | NA | [10] |
| FBXW4 | BLCA | Supp | NA | [22] |
| HAND2 | BLCA | Supp | NA | [32] |
| HAND2 | UCEC | Supp | NA | [10] |
| FOXF1 | BLCA | Supp | NA | New |
| FGFR2+FBXW4 | BLCA | Supp | Coop | New |
| FGFR2+HAND2 | UCEC | Supp | Ant | New |

Table S19: Summary of reported findings. Column Key: Genes = the genes in the specified relationship, Cancer = BLCA or UCEC, Effect = Oncogenic or Suppressor, Effect Type = NA (single gene), Antagonistic or Cooperative, Status = Known or Unknown, Citation = reference utilized in the manuscript to reference a known effect

[5] Ricardo J. G. B. Campello, Davoud Moulavi, and Joerg Sander. Density-Based Clustering Based on Hierarchical Density Estimates. In Jian Pei, Vincent S. Tseng, Longbing Cao, Hiroshi Motoda, and Guandong Xu, editors, *Advances in Knowledge Discovery and Data Mining*,

Lecture Notes in Computer Science, pages 160–172, Berlin, Heidelberg, 2013. Springer.

[6] Hung-Chia Chen, Ralph L Kodell, Kuang Fu Cheng, and James J Chen. Assessment of performance of survival prediction models for cancer prognosis. *BMC medical research methodology*, 12(1):102, 2012.

[7] Antonio Colaprico, Tiago C Silva, Catharina Olsen, Luciano Garofano, Claudia Cava, Davide Garolini, Thais S Sabedot, Tathiane M Malta, Stefano M Pagnotta, Isabella Castiglioni, et al. Tcgabiolinks: an r/bioconductor package for integrative analysis of tcga data. *Nucleic acids research*, 44(8):e71–e71, 2016.

[8] Gene Ontology Consortium. The gene ontology resource: 20 years and still going strong. *Nucleic acids research*, 47(D1):D330–D338, 2019.

[15] John P. Klein, Hans C. Van Houwelingen, Joseph G. Ibrahim, and Thomas H. Scheike. *Handbook of survival analysis.* CRC Press, 2016.

[23] Leland McInnes, John Healy, and James Melville. Umap: Uniform manifold approximation and projection for dimension reduction. *arXiv preprint arXiv:1802.03426*, 2018.

[24] Amanda J Mikels and Roel Nusse. Purified wnt5a protein activates or inhibits $\beta$-catenin–tcf signaling depending on receptor context. *PLoS biology*, 4(4), 2006.

[25] Virginia Murillo-Garzón, Irantzu Gorroño-Etxebarria, Malin Åkerfelt, Mikael Christer Puustinen, Lea Sistonen, Matthias Nees, James Carton, Jonathan Waxman, and Robert M Kypta. Frizzled-8 integrates wnt-11

and transforming growth factor-$\beta$ signaling in prostate cancer. *Nature communications*, 9(1):1–16, 2018.

[26] Manfred Opper and Ole Winther. Gaussian processes for classification: Mean-field algorithms. *Neural computation*, 12(11):2655–2684, 2000.

[27] Carl Edward Rasmussen and Christopher KI Williams. Gaussian processes for machine learning, vol 14, no 2, 2006.
