## Supplemental Figures for "Modeling the effects of EMT-immune dynamics on epithelial cancer progression"

### Supplementary Figures for Modeling the competing effects of the immune system and EMT on epithelial cancers

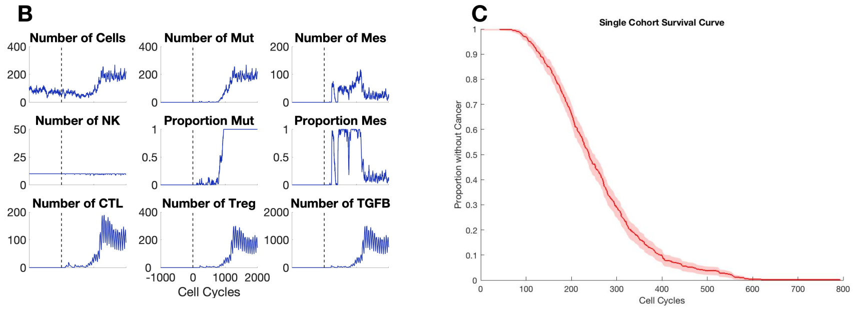


Figure S1: A. Single patient trajectory without immune cells targeting cells with pathways mutations. Compare to Fig. 1B. B. Sample cohort survival curve without immune cells targeting cells with pathways mutations. Compare to Fig. 1C.

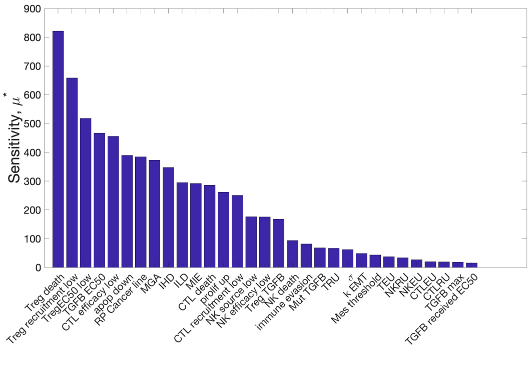


Figure S2: Morris-OAT global sensitivity without immune cells targeting cells with pathways mutations. Compare to Fig. 2.

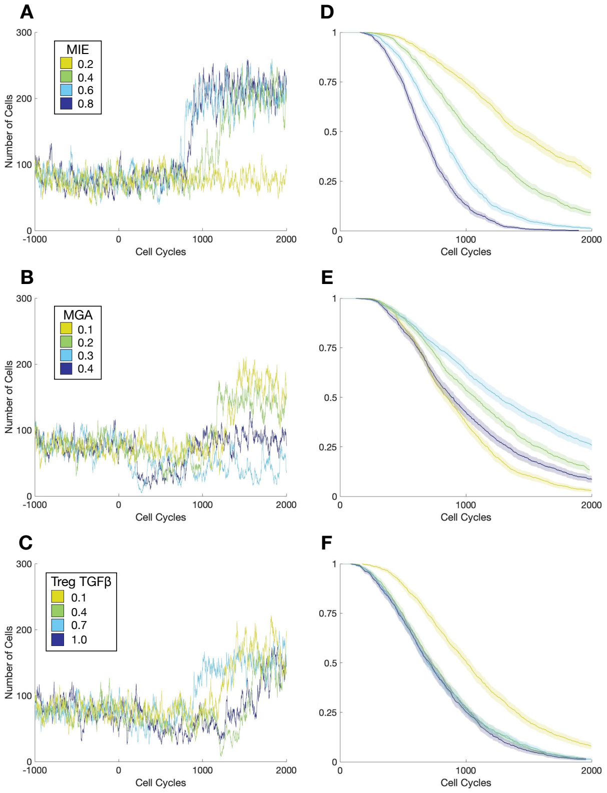


Figure S3: Effects of mesenchymal tumor cell properties on the Time to Invasion without immune cells targeting cells with pathways mutations. Compare to Fig. 3.

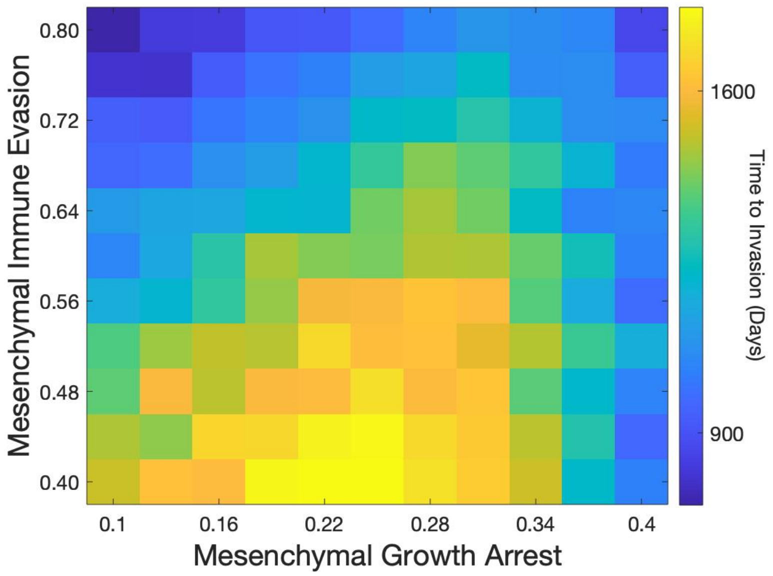


Figure S4: Summary of mesenchymal tumor cell properties on the Time to Invasion without immune cells targeting cells with pathways mutations. Compare to Fig. 4E.

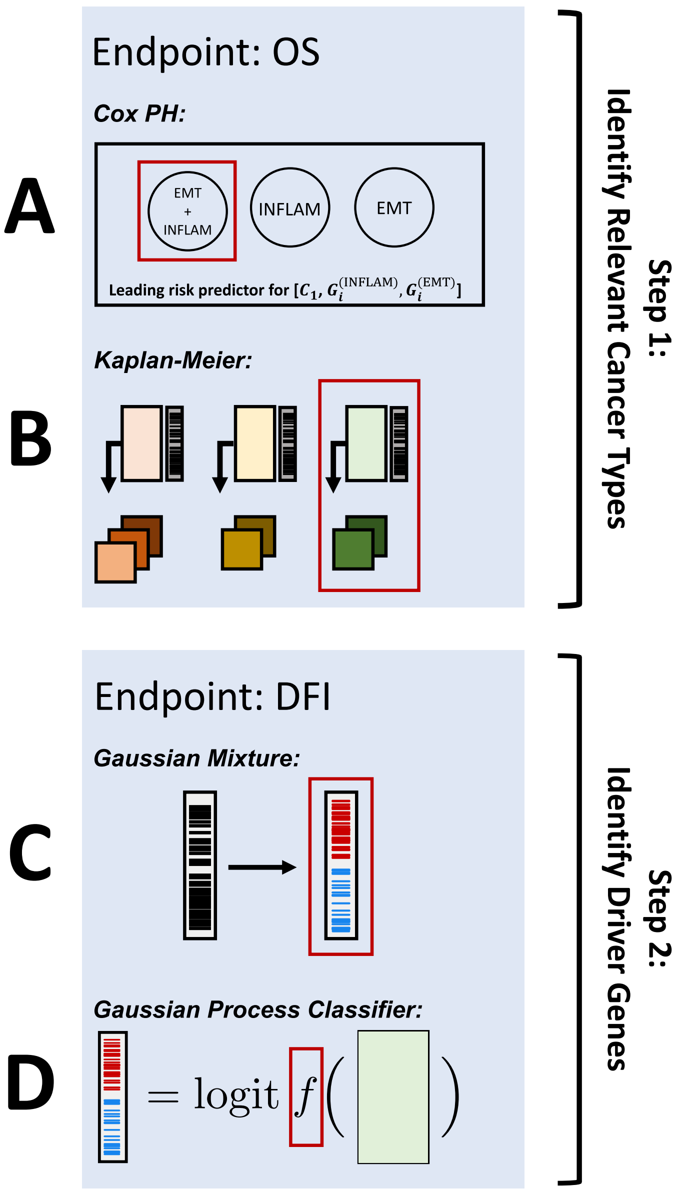


Figure S5: Schematic of overall survival (OS) and disease-free interval (DFI) analysis on TCGA data. A-B (Step 1): Find cancer types in which EMT and inflammation act synergistically on overall survival. C-D (Step 2): For cancer types identified in Step 1, identify proliferation pathways that regulate the invasiveness of these cancers. A) For each cancer type, identify pairs of inflammation and EMT gene sets where the UMAP projection over their union has a higher (magnitude) hazard ratio than either of its constituents in a three-predictor Cox-PH model with OS response in that cancer type. B) Identify cancer types, for which unsupervised DBSCAN clustering over the 1D UMAP projection of one or more EMT/Inflam union sets yields clusters whose KM survival curves are different. C) Impute the DFI-high/low class based on a two-component Gaussian mixture model of the published disease-free interval time in days. D) Identify relationships between proliferation pathways and tumor invasiveness using a GP classifier trained on the computed DFI-class from (C) and mRNA sequencing for the list of proliferation genes.

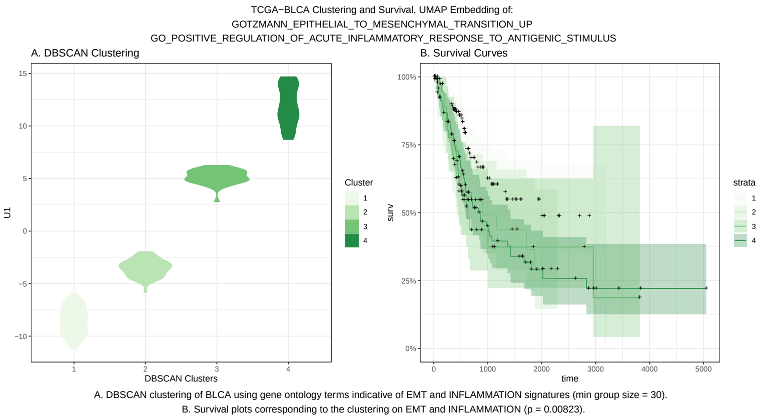


Figure S6: DBSCAN clusters for combined EMT+INFLAM embedding of BLCA patients and corresponding Kaplan-Meier (KM) model.

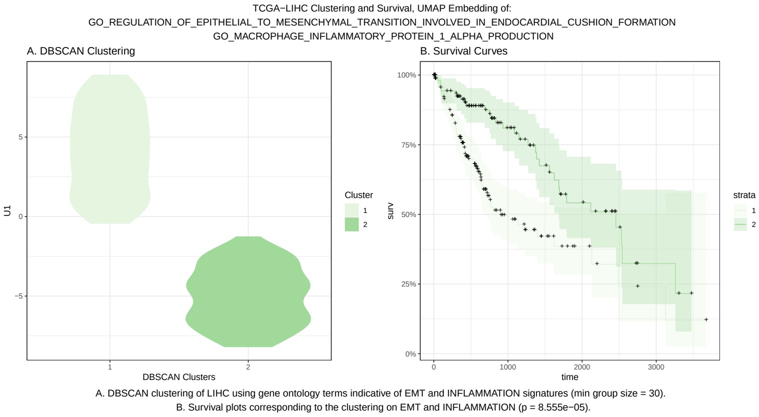


Figure S7: DBSCAN clusters for combined EMT+INFLAM embedding of LIHC patients and corresponding Kaplan-Meier (KM) model.

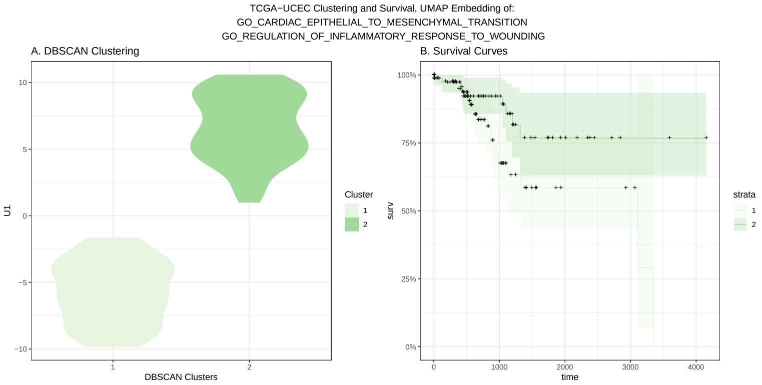


A. DBSCAN clustering of UCEC using gene ontology terms indicative of EMT and INFLAMMATION signatures (min group size = 30).
B. Survival plots corresponding to the clustering on EMT and INFLAMMATION (p = 0.02372).

Figure S8: DBSCAN clusters for combined EMT+INFLAM embedding of UCEC patients and corresponding Kaplan-Meier (KM) model.

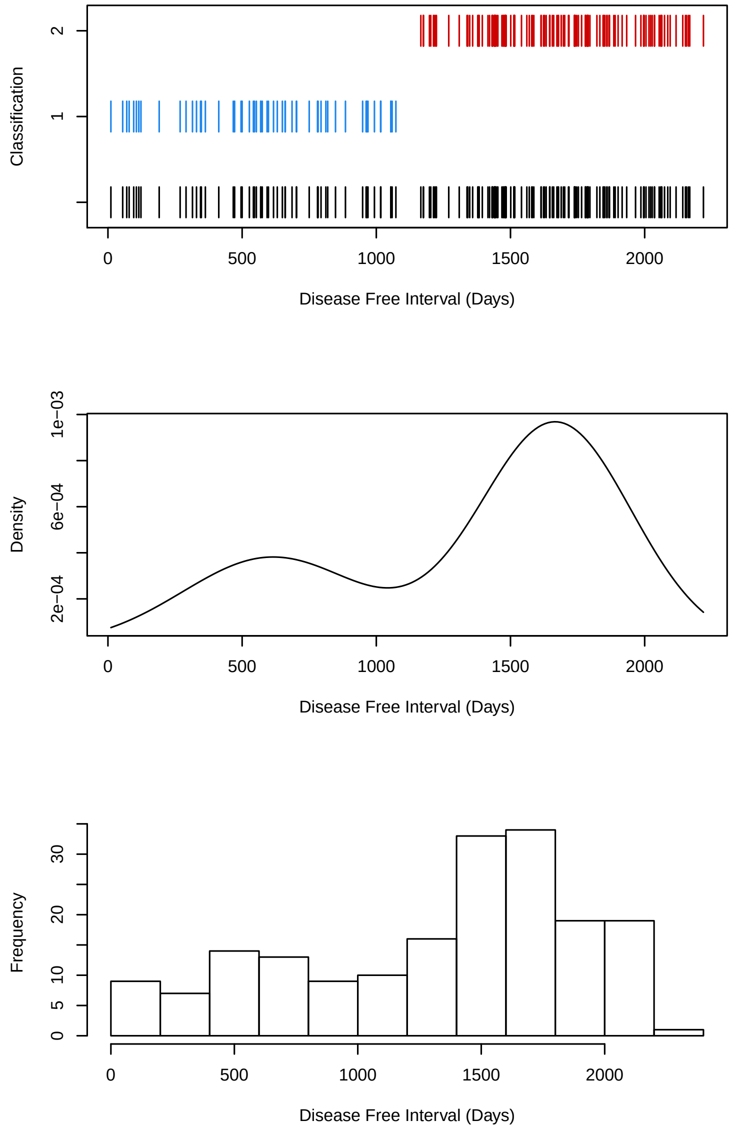


Figure S9: A. Classification of BLCA patients in short DFI (blue) and long DFI (red). B. 2-component Gaussian mixture density corresponding to the above classification. C. Histogram of DFI for patients.

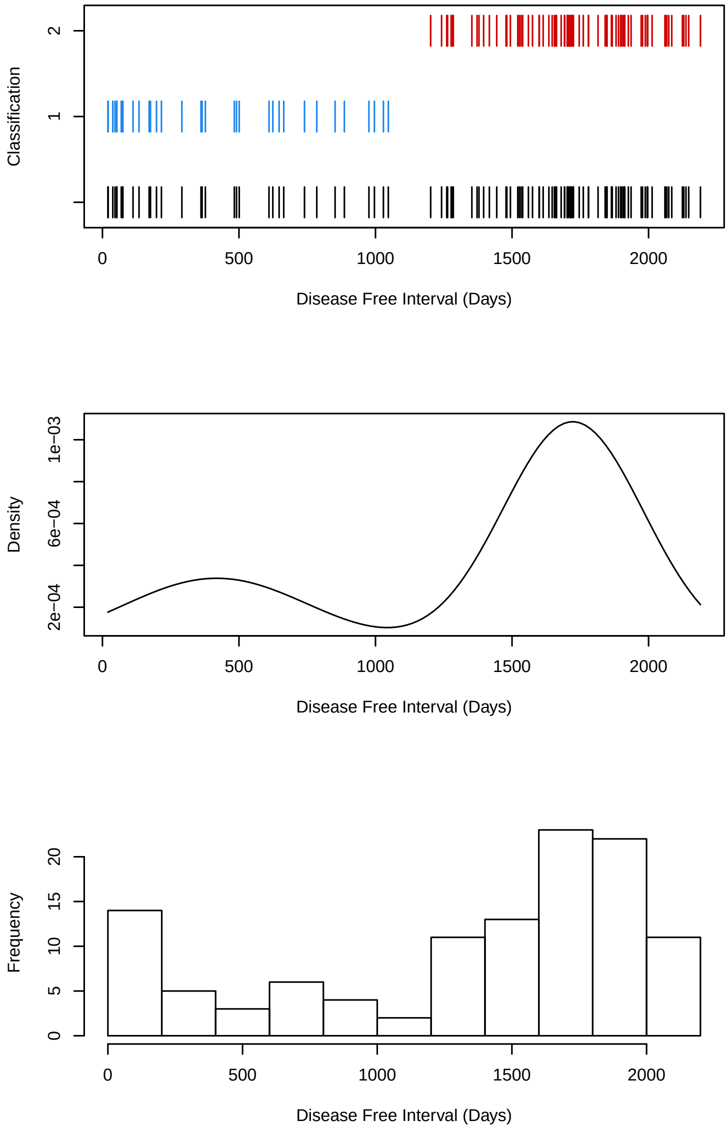


Figure S10: A. Classification of UCEC patients in short DFI (blue) and long DFI (red). B. 2-component Gaussian mixture density corresponding to the above classification. C. Histogram of DFI for patients.

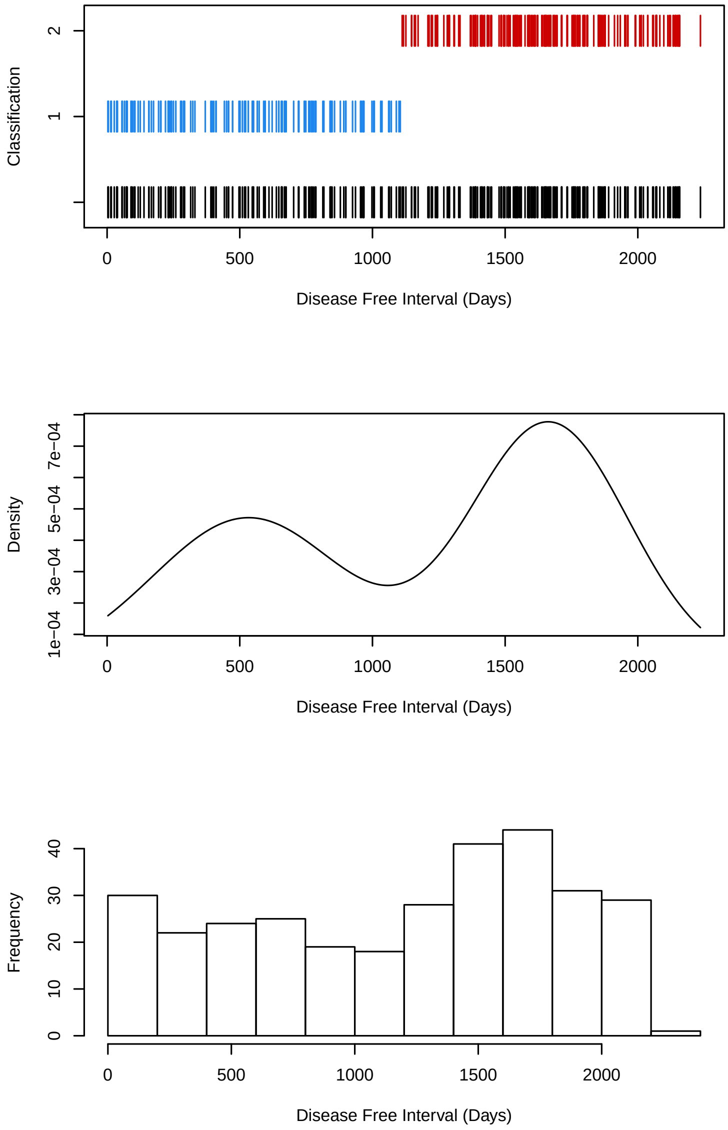


Figure S11: A. Classification of LIHC patients in short DFI (blue) and long DFI (red). B. 2-component Gaussian mixture density corresponding to the above classification. C. Histogram of DFI for patients.
